# Long-range connections mirror and link microarchitectural and cognitive hierarchies in the human brain

**DOI:** 10.1101/2021.10.25.465692

**Authors:** Yezhou Wang, Jessica Royer, Bo-yong Park, Reinder Vos de Wael, Sara Larivière, Shahin Tavakol, Raul Rodriguez-Cruces, Casey Paquola, Seok-Jun Hong, Daniel S. Margulies, Jonathan Smallwood, Sofie L. Valk, Alan C. Evans, Boris C. Bernhardt

## Abstract

Core features of higher-order cognition are hypothesized to be implemented via distributed cortical networks that are linked via long-range connections. However, these connections are biologically expensive, and it is unknown how the computational advantages long-range connections provide overcome the associated wiring costs. Our study investigated this question by exploring the relationship between long-range functional connections and local cortical microarchitecture. Specifically, our work *(i)* profiled distant cortical connectivity using resting-state fMRI and cortico-cortical geodesic distance mapping, *(ii)* assessed how long-range connections reflect local brain microarchitecture, and *(iii)* studied the microarchitectural similarity of regions connected through long-range connections. Analysis of two independent datasets indicated that sensory and motor areas had more clustered short-range connectivity patterns, while transmodal association cortices, including regions of the default mode network, were characterized by distributed, long-range connections. Confirmatory meta-analysis suggested that this topographical difference mirrored a shift in cognitive function, from perception/action towards emotional and social cognitive processing. Analysis of myelin-sensitive *in vivo* MRI in the same participants as well as *post mortem* histology and gene expression established that gradients in functional connectivity distance are paralleled by those present in cortical microarchitecture. Moreover, long-range connections were found to link together spatially remote regions of association cortex with an unexpectedly similar microarchitecture. These findings provide novel insights into how the organization of distributed functional networks in transmodal association cortex contribute to cognition, because they suggest that long-range connections link together distant islands of association cortex with similar microstructural features.

## Introduction

In the mammalian brain, the distance between cortical regions largely determines the extent to which they are inter-connected ^1,2^. Although most cortico-cortical connections are local and short-range ^3^, it is hypothesized that long-range connectivity patterns likely play a key role in organizing human brain structure and function ^4^. Long-range connections may help integrate distributed functional communities facilitating more complex computation; thus, their increased wiring cost is balanced by their benefits to network efficiency in inter-network communication ^5-9^. Long-range connections, however, are neither evenly nor randomly distributed across the cortical mantle, but appear concentrated in transmodal association cortex, networks that engage in increasingly abstract and self-generated cognition ^10-13^. Conversely, long-range connections are less frequently found in sensory and motor regions that interact more closely with the here and now ^10,14-16^. Despite the widely held view that long-range connectivity holds functional importance to human brain organization, the specific layout and neural underpinnings of long-range connections remain incompletely characterized.

The study of human brain connectivity has increasingly benefitted from the analysis of coordinated brain signals during *resting-state* functional magnetic resonance imaging (fMRI) acquisitions ^17,18^. Resting-state fMRI (rs-fMRI) probes multiple functional networks within a single acquisition ^18-22^, and rs-fMRI networks have been shown to be both reproducible and reliable ^23^. Prior research has underscored the value of rs-fMRI in profiling connectivity patterns of specific areas ^24-26^ and of delineating large-scale networks, which often correspond to systems engaged during cognitive states ^27^. To understand distance effects on cortical connectivity, prior studies combined rs-fMRI analysis with cortico-cortical distance profiling ^4,10^, suggesting that sensory/motor networks have a locally clustered connectivity profile while transmodal association cortices show rather long-range connections. Here, we further extend these approaches and capture functional connectivity in an anatomically grounded way, by analytically differentiating regions embedded within long-*vs* short-range functional networks.

Converging evidence has indicated that cortical function generally follows similar spatial axes as microstructural variations ^28,29^. Spatial gradients that differentiate sensory and motor systems from transmodal association cortices have been reported at the level of cortical microstructure captured using myelin-sensitive *in vivo* MRI as well as *post mortem* 3D histology ^30^. Such sensory-fugal spatial trends have also been indicated when assessing cortical gene expression, including recent data based on *post mortem* microarray transcriptomics from the Allen Brain Institute showing regionally varying cortical microcircuit organization ^31^. Notably, these gradients appear to align with gradients obtained from functional connectivity analysis differentiating sensory/motor systems from transmodal networks in term of intrinsic connectivity patterns ^32,33^, suggesting overall correspondence between functional hierarchies and microcircuit patterns ^28,34,35^. Here, we built on this emerging literature, by performing systematic spatial association analyses between functional connectivity distance topographies as well as cortical microarchitecture derived from myelin-sensitive *in vivo* MRI in the same participants, as well as *post mortem* 3D histology and transcriptomics.

In addition to examining spatial covariation of function and structure across the cortical mantle, combined structure-function assessment can clarify the role of long-range connectivity in the global cortical landscape. In general, cortical regions that are closer to one another are more likely to be connected and will have a more similar microstructural context. Conversely, there is generally an overall reduction in the microstructural similarity and functional connectivity of different areas with increasing spatial distance between them ^29,34,36^. However, the human brain seems to also incorporate notable exceptions to this rule ^16,36,37^. For example, transmodal association cortices show an increased clustering of (otherwise relatively rare) long-range cortico-cortical connections, and distributed association networks also more likely to be connected than expected based on their sometimes-large distance. By studying the similarity of cortical functional networks in terms of their microstructure and gene expression profiles, we furthermore aimed at testing whether long-range connectivity patterns serve a special role in an expanded cortical surface, allowing to connect microstructurally similar cortices across a large cortical distance.

The current work mapped cortical topographies of functional connectivity distance in the human brain and examined associations to underlying cortical microstructure. We profiled the embedding of each cortical region within long-*vs* short-range functional networks based on a combination of rs-fMRI connectivity and cortex-wide geodesic distance mapping ^4,38^. In a first set of analyses, we contextualized the topography of long-range connections with respect to other established motifs of brain function as well as meta-analytical cognitive ontologies. We furthermore examined associations to cortical microarchitecture, combining analysis of myelin-sensitive *in vivo* MRI, *post mortem* histology, and *post mortem* gene expression. To further understand the role that long-range connections play in large-scale brain function in an expanded human cortex, we assessed the relationship between spatial patterns of functional connectivity distance and the similarity of connected areas in terms of cortical microarchitecture. Several sensitivity analyses assessed robustness of our findings with respect to different analytic choices, and we replicated all *in vivo* findings in an independent dataset.

## Results

Our main analyses were based on a sample of 50 healthy young adults (29.82 ± 5.73 years, 21 females) evaluated in our institute who had high definition functional and microstructural *in vivo* MRI data available ^39^. The replication analysis was based on an independent dataset of 200 unrelated healthy young adults from the Human Connectome Project (HCP) dataset ^40^ (28.68 ± 3.69 years, 119 females).

### Gradients of functional connectivity distance

We profiled a region’s functional connectivity distance by mapping its embedding into functional connectivity subgraphs with a specific spatial distance. Specifically, we calculated for each region the correlation between its connectivity distance, which is the average geodesic distance of its functionally connected brain regions, and functional connectivity (see *Methods*). We observed highest functional connectivity distance in transmodal association networks and lowest functional connectivity distance in sensory/motor cortices (**Fig. 1A**).

**Fig. 1.**
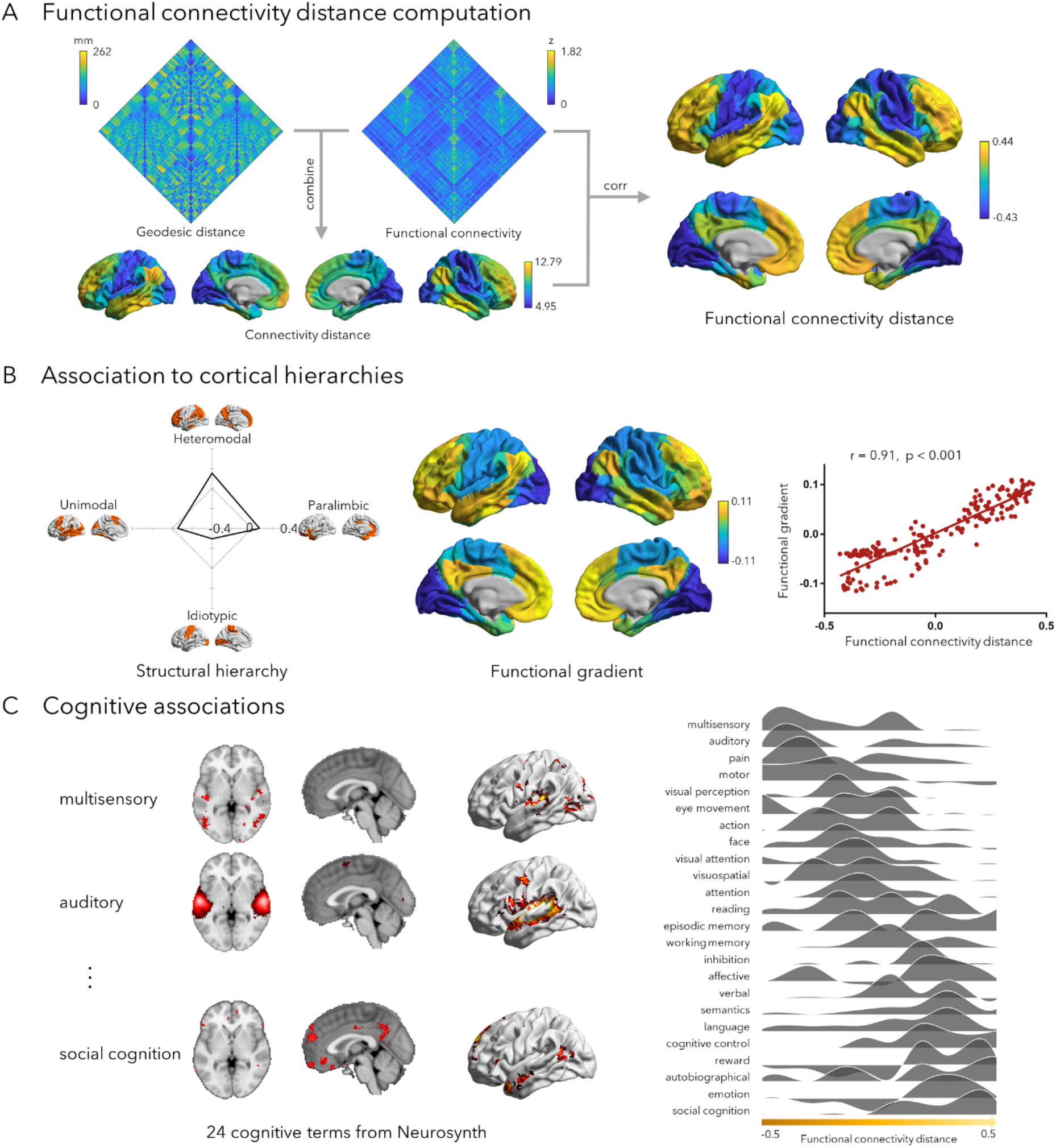
The topography of functional connectivity distance in the human brain. **(A)** Connectivity distance was calculated from the combination of cortex-wide geodesic distance measures and functional connectivity analysis, and tapped into the embedding of a cortical region into short-vs long-range functional subnetworks. **(B)** Stratification relative to a model of the primate cortical hierarchy ^29^, and a data driven gradient of rs-fMRI connectivity. **(C)** Cognitive associations, based on confirmatory task fMRI meta-analysis using Neurosynth ^42^.

When stratifying functional connectivity distance according to a model of the primate cortical hierarchy and laminar differentiation ^29^, we identified peak values in heteromodal and paralimbic cortices, the transmodal apex of the sensory-fugal hierarchy, while lowest values were observed in primary cortices (**Fig. 1B**). We also examined associations to the principal functional gradient, an approximation of the cortical hierarchy derived non-linear dimensionality reduction applied to rs-fMRI connectivity data ^43^. Controlling for spatial autocorrelation, the principal gradient was found to closely resemble the functional connectivity distance topography (r = 0.91, p_spin_ < 0.001). Similar findings were observed when stratifying functional connectivity distance according to intrinsic functional communities ^41^, (***SI Appendix*, Fig. S1**). Moreover, we also observed a moderately positive correlation between functional connectivity distance and the function participation coefficient (r = 0.30, p_spin_ = 0.003), a graph theoretical metric of diverse/integrative connectivity of a region ^44-46^, suggesting that areas with longer connectivity distance also showed more integrative connectivity profiles.

Confirmatory meta-analysis using the Neurosynth ^42^ database of previous task-fMRI studies estimated spatial associations to activation patterns across a set of diverse cognitive terms ^33^. The resulting spatial associations were stratified from low to high functional connectivity distance (**Fig. 1C**). Terms associated with long-range connectivity involved social/emotional functions, such as ‘social cognition’ and ‘emotion’ while short-range connections were more common in processes related to sensory functions, such as ‘multisensory’ and ‘auditory’.

We evaluated robustness of the functional connectivity distance findings against variations in preprocessing and analysis choices (*i*.*e*., functional connectome thresholding and use of global mean signal regression during preprocessing), and obtained similar results **(*SI Appendix*, Fig. S2)**. Functional findings were also relatively stable at the individual participant level **(*SI Appendix*, Fig. S3)**. Moreover, functional findings were similar when analyzing the independent HCP dataset (***SI Appendix*, Fig. S4**).

### Associations to local cortical microarchitecture

We next examined associations to cortical microarchitecture. Firstly, we sampled intracortical *in vivo* quantitative T1 (qT1) relaxometry MRI profiles ^*47,48*^ in all cortical parcels in the same participants. We then cross-correlated qT1 intensity profiles between parcels to generate a microstructural similarity matrix (**Fig. 2A**), and generated a cortex-wide microstructural gradient ^28,43^, running from primary towards paralimbic areas. Functional connectivity distance correlated strongly with this gradient (r = 0.75, p_spin_ < 0.001; **Fig. 2A**). Correlations remained stable at the individual level (***SI Appendix*, Fig. S5A**). We confirmed that structure-function relations were similar in the HCP dataset (***SI Appendix*, Fig. S6**), which uses T1w/T2w imaging ratio as an alternative microstructural index ^49^.

**Fig. 2.**
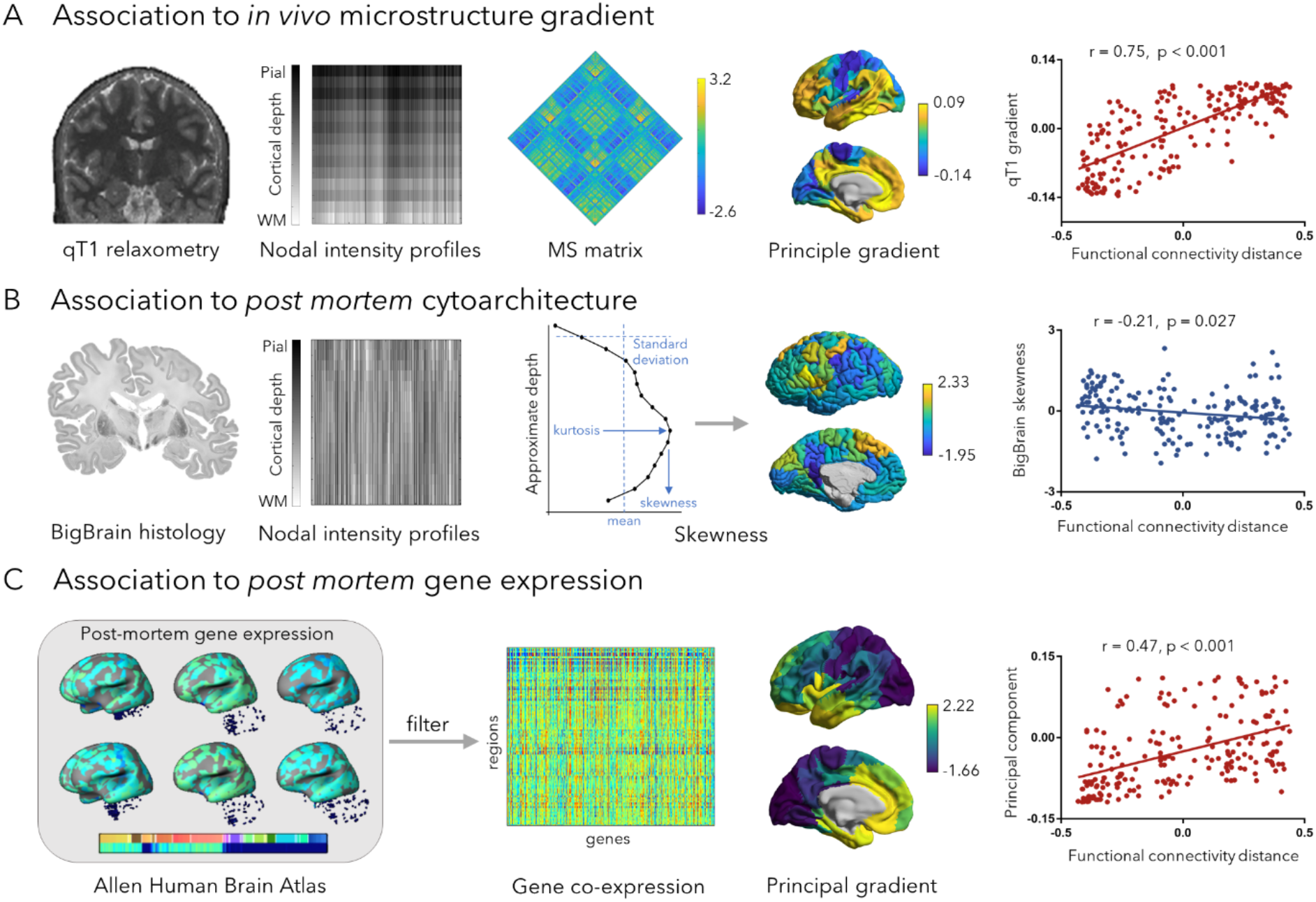
Associations to cortical microarchitecture. **(A)** Association with the *in vivo* microstructural gradient based on quantitative T1 (qT1) relaxometry obtained in the same participants. **(B)** Association with *post mortem* histological measures (intracortical profile skewness) derived from the BigBrain dataset ^50^. (C) Association with *post mortem* gene expression data derived from the Allen Human Brain Atlas ^53^. Gene expression data was filtered across 6 donors, and a gene co-expression matrix was constructed followed by principal component analysis. *Abbreviation*: MS, microstructural similarity.

In a separate analysis, we sampled intracortical intensity profiles from BigBrain^28,50^, a 3D *post mortem* histological reconstruction of a human brain. We estimated skewness measures ^51^ of intensity profiles to represent microstructural differentiation ^29,52^, and obtained significant correlations with functional connectivity distance (skewness: r = -0.21, p_spin_ = 0.027; **Fig. 2B**). We also assessed skewness of qT1 intensity profile and obtained significant correlations with functional connectivity distance at group and individual level (r = -0.72, p_spin_ < 0.001; ***SI Appendix*, Fig. S5B**).

We leveraged the Allen Human Brain Atlas ^53^ to assess transcriptomic correlates of connectivity distance. We selected only genes consistently expressed across the six donors (r > 0.5; 1050/20647 (5.09%) genes (**Fig. 2C, *SI Appendix*, Table S1**). Based on those 1050 genes, we identified the first principal component of gene expression (explaining 71.58% of co-expression variance) and found that this component correlated to the spatial pattern of long-range connections (**Fig. 3B**, r = 0.47, p_spin_ < 0.001). We then performed cell-type specific gene enrichment ^54,55^ analysis and calculated overlap ratios in gene expression between the 508 positively distance-related genes (***SI Appendix*, Table S2**) and cell-type specific genes using 1,000 permutation tests. These genes showed a significant overlap with genes expressed in excitatory and inhibitory neurons in both supragranular and infragranular layers, as well as astrocytes and oligodendrocyte progenitor cells (FDR < 0.05; ***SI Appendix*, Fig. S7**). Therefore, functional connectivity distance was found paralleling cortical microarchitectural gradients derived from myeloarchitecture, cytoarchitecture and gene expression.

**Fig. 3.**
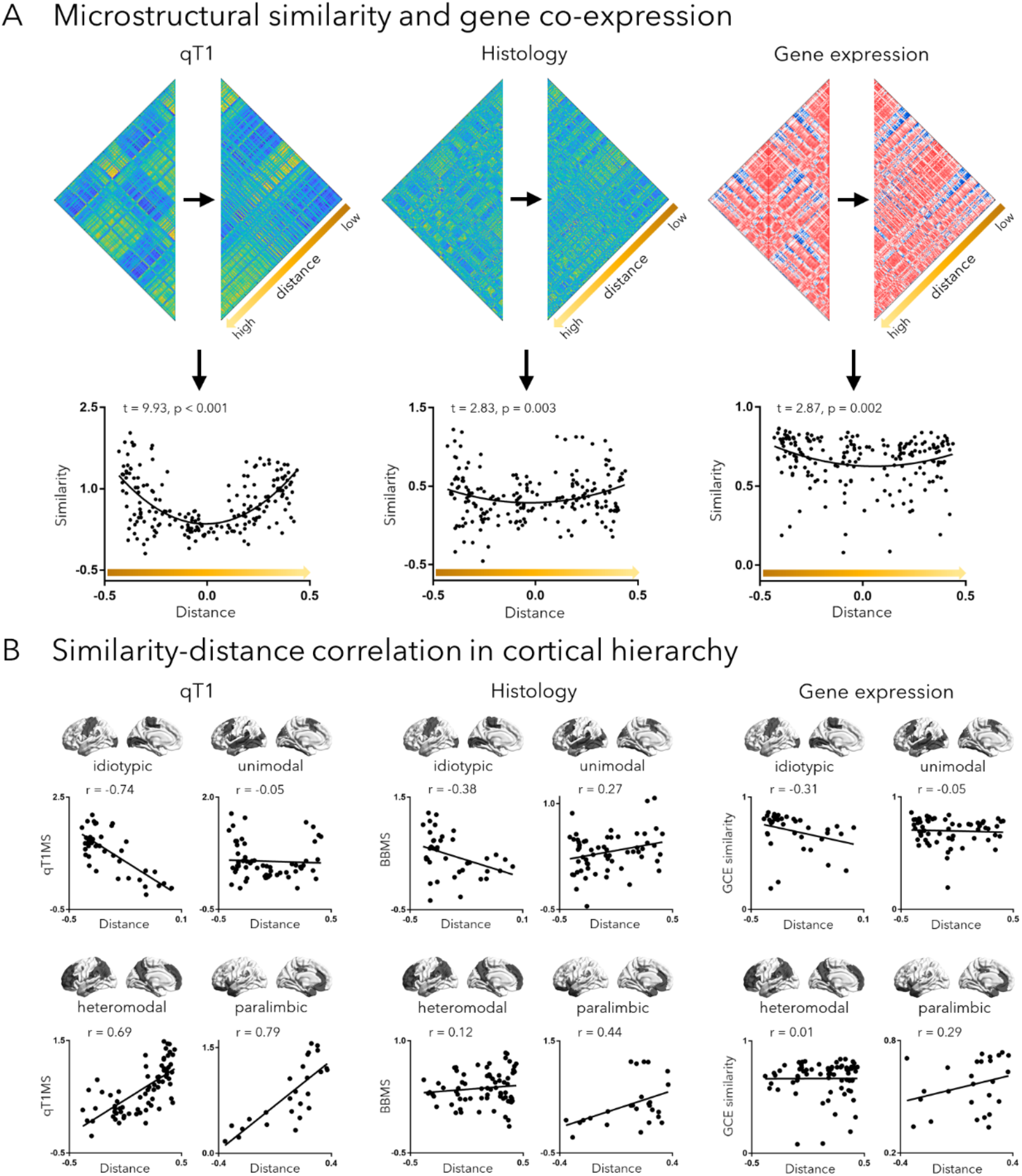
Long-range connections bypass distance dependent wiring in the human brain. (A): The microstructural similarity (MS) matrices (derived from qT1 and BigBrain) and gene co-expression matrix were sorted by functional connectivity distance and averaged in the predefined functional connectivity configuration to generate the microstructural and genetic similarity map. These maps were then fitted to functional connectivity distance using quadratic function. (B): The correlation between microstructural/genetic similarity and functional connectivity distance was further examined in cortical hierarchy. *Abbreviation*: qT1MS, microstructural similarity derived from qT1; BBMS, microstructural similarity derived from BigBrain; GCE, gene co-expression.

### How long-range connectivity departs from distance-dependent cortical wiring rules

In the human brain, microarchitectural similarity of cortical regions generally declines with increasing spatial distance ^34,36^. Here, we verified this relationship and assessed the role of long-range connections in such distance-dependent microarchitectural associations. We quantified microstructural similarity as above using qT1, BigBrain, and gene co-expression of cortico-cortical functional networks, and reordered similarity according to functional connectivity distance (**Fig. 3A**). Interestingly, we observed a quadradic relationship between microstructural similarity and functional connectivity distance for qT1 (t=9.93, p<0.001), BigBrain (t=2.83, p=0.003), and gene expression similarity (t=2.87, p=0.002). In all cases, a quadradic model provided a better model fit than a linear model (F>8.03, p<0.001), suggesting that regions with high connectivity distance express and unexpectedly high microarchitectural similarity given their distance (***SI Appendix*, Fig. S8**). An overarching mixed effects model, aggregating qT1, BigBrain, and gene expression confirmed a strong quadratic effect (t=9.04, p<0.001). When assessing distance effects stratified for different levels of the cortical hierarchy levels, negative effects of distance on microstructural/genetic similarity were found in primary and unimodal association networks, while transmodal areas (*i*.*e*., heteromodal and paralimbic networks) showed positive relations (**Fig. 3B**). Similar findings were seen for different intrinsic functional communities (***SI Appendix*, Fig. S9**). Moreover, findings were robust in the HCP dataset **(*SI Appendix*, Fig. S10**).

### Data Availability

Data from the locally studied dataset are openly available on CONP (https://portal.conp.ca/dataset?id=projects/mica-mics); processing pipelines are openly available on http://github.com/MICA-MNI/micapipe documented on http://micapipe.readthedocs.io.

## Discussion

Long-distance connections are likely key for distributed human brain function, and it is widely assumed that the associated high wiring costs are offset by gains in global communication efficiency ^5-9,56,57^. The present study integrated rs-fMRI connectivity analysis and cortex-wide geodesic distance mapping, and differentiated transmodal regions embedded within long-range networks from sensory and motor networks embedded in short-range subgraphs. Increases in connectivity distance were found to reflect increasingly integrative connectivity patterns ^28,34,58^ and clustered towards the apex of a sensory-transmodal gradient of intrinsic brain function ^28,32,33^. Confirmatory meta-analysis of previous task-fMRI findings based on Neurosynth^42^ demonstrated that these regions are engaged in more affective and higher-order social processes. In contrast, regions involved in sensory and motor functions were mainly characterized by more localized, short-range cortico-cortical connections. Capitalizing on *in vivo* myelin-sensitive MRI obtained in the same participants, as well as *post mortem* 3D histology and transcriptomics datasets ^50,53^, we determined that the topography of functional connectivity distance converged with sensory-fugal gradients in cortical microarchitecture. Critically, long-range connections were found link spatially remote islands of transmodal association cortices with high microarchitectural similarity. Establishing topographic associations between long-range connections and cortical microarchitecture showed that long-range connections partially counteract the overall distance-dependent decline in inter-areal similarity, thus providing a possible substrate for integrative and parallel function in human association cortex.

Our systematic analysis of short-*vs* long-range functional network embedding complements prior studies that have explored cortical connectivity distance variations in human and non-human brains. Using a discrete Euclidean distance thresholding (set at 14 mm) of resting-state fMRI connectivity patterns, a prior study categorized areas into those with local *vs* distant connections ^10^. Parallel graph-theoretical assessment revealed that regions with short-range connections more frequently contributed to localized network clustering, while long-range connections ensure global efficiency increases ^10^. Follow-up studies in humans and non-human primates combined rs-fMRI and surface-based geodesic distance mapping and suggested dissociations between sensory and transmodal regions in terms of wiring distance ^4,59^. Here, we expanded these previous methods by using a formulation that integrates the connectivity distance of a given region with functional connectivity profiles of immediately connected neighbors of that region, to index the distance-dependent subgraph embedding of a given area. This approach markedly segregated transmodal areas from unimodal and sensory systems, in line with work suggesting that midline regions in paralimbic and default mode networks have a high-density and long-range embedding, an architecture also referred to as the *rich-club* ^60^. In our study, we confirmed that long-range connectivity directly contributes to integrative brain function by showing spatial correlations between connectivity distance and the participation coefficient, a graph-theoretical index of the connectional diversity of a given region ^44^. Diverse connectivity profiles likely play a role in network stability, along with the maintenance of whole-brain communication ^61^. We also noted robust correlations to the principal functional connectivity gradient ^33^, a data-driven approximation of sensory-transmodal hierarchy in the primate brain ^28,29,33^. Our results, thus, support that functional connectivity distance variations reflect established motifs of macroscale cortical organization. Confirmatory meta-analysis based on neurosynth ^42^ revealed that connectivity distance variations mirrored, in part, putative cognitive hierarchies, running from sensory and motor processes towards higher-order affective and socio-cognitive functions ^28^. Uni- and heteromodal association cortices underwent a disproportionate enlargement during human evolution ^62,63^, and long-range connections may, in turn, ensure their continued ability to communicate across a wide cortical territory. This may, overall, provide increased ability to integrate diverse information, and thus allowed for a greater degree of cognitive flexibility and adaptability of these networks.

A mounting literature emphasizes that cortical functional organization likely reflects variations in cortical microarchitecture ^32,64-67^. Using myelin-sensitive *in vivo* MRI in the same participants ^68-71^, we could show that the sensory-fugal connectivity distance patterns follow gradients that distinguish sensory and motor regions from transmodal cortices on the basis of intracortical microstructure ^33,34,66,72^. The specific microstructural context of transmodal *vs* sensory/motor cortices may show different neurodevelopmental trajectories and diverging potential for experience-driven plasticity ^73^, a finding likely related to regional variations in activity-dependent myelination ^74^ and synaptic pruning ^29,75^. Associations to intracortical microstructure, in particular cytoarchitectonic measures of laminar differentiation, were also seen when examining associations on a *post mortem* 3D reconstruction of a human brain, the BigBrain dataset ^50^. Our approach furthermore showed transcriptomic co-variations with shifts in functional connectivity distance ^53^. A congruence was globally seen when studying associations to the principal component of cortical gene co-expression, echoing prior findings on transcriptomic substrates of functional imaging measures ^31,76,77^. Paralleling the myelo- and cytoarchitectural underpinnings of neuronal function, the transcriptional and functional architecture of human cortex likely share a common axis, where gradients of microscale properties ultimately contribute to macroscale functional specialization ^66^. Gene enrichment analysis could identify associations to genes expressed in excitatory and inhibitory neurons in both supragranular and infragranular compartments, together with astrocytes. Overall, this finding points to a rather diverse microcircuit layout of transmodal regions harboring long-range connections. A recent study reported genes enriched in supragranular cortical layers in humans relative to mice ^78^, molecular innovations that may have contributed to the evolution of long-range cortico-cortical projections. Another study systematically identified long-range projecting markers by aligning temporal variation in gene expression and structure, reporting a set of genes identifying long-range projecting neurons in both upper and deeper layers ^79^. Neuronal as well as non-neuronal contributions to microcircuit organization, for example through astrocytes may play important roles in neuronal activity ^80,81^ and plasticity ^82^ at both excitatory and inhibitory synapses ^83,84^. Notably, by acting through transmitter-dependent and -independent mechanisms ^85-90^, astrocytes cells can serve as modulators of short-as well as long-range cortico-cortical communication ^91^.

Generative connectome models seek to identify the rules governing cortical wiring ^92^. Studies overall show that a combination of geometric constraints, including a distance penalty, as well as a homophilic wiring mechanism can reproduce key topological characteristics of the human connectome ^92,93^. Distance-dependence implicates that proximal regions are more likely to be connected ^94^. Similarly, findings in many species indicate that areas with similar microstructural and neurobiological features are more likely to be wired to one another ^36,37,95,96^. Our work adds to this literature, suggesting an intriguing interplay of both distance dependence and homophilic wiring, underscored by the observed u-shaped association between microarchitectural similarity and connectivity distance. This confirms, on the one hand, the overall notion that adjacent regions with short-range connections have similar gene expression and microstructure, potentially underpinning their segregative functional roles. On the other hand, it also shows that transmodal cortices that participate in the long-range functional subnetworks have an unexpected increase in their microarchitectural similarity. The rare long-range connections that are mainly present in transmodal association cortices, thus, appear to allow for distant communication of microstructurally and transcriptionally similar territories, bypassing the general distance-dependent decline in homophily of brain wiring. Long-range connections, thus, likely allow for integrative communication across remote cortical regions ^60,97^. Thereby, communication pathways formed by long connections may exhibit redundancies that promote network robustness and complexity, and potentially enable parallel information processing across different cortical zones ^56^.

A battery of sensitivity analyses confirmed that that our findings were relatively robust across different analysis parameter choices, and we replicated our main findings in an independent data set and showed consistency at both the group- and single participant-level. Overall, these results thus provide evidence for a sensory-fugal topography of functional connectivity distance variations and show spatially covarying microarchitectural axes. Long-range connections may allow for integration of spatially remote, yet microarchitecturally similar areas of association cortex. These results suggest that long-range connections may facilitate integrative communication in a massively expanded cerebral cortex, and thus potentially contribute to parallel and higher order information processing that is central to human cognition.

## Materials and Methods

A detailed description of the Subjects inclusions, image processing, and analysis methodology can be found in the ***SI Appendix, in the Supplemental Materials and Methods***. In brief, we studied 50 unrelated healthy adults (age: 29.82±5.73 years, 21 females) ^39^. All participants had quality-controlled multimodal MRI data available (see below). The Ethics Committee of the Montreal Neurological Institute and Hospital approved the study. Written informed consent was obtained from all participants.

Based on rs-fMRI, we constructed functional connectomes on 200 functionally defined cortical parcels ^98^. We combined connectivity distance and functional connections to generate functional connectivity distance, which highlighted the embedding of each cortical region within long-vs short-range functional networks. We stratified functional connectivity distance according to intrinsic functional communities ^41^ and a model of primate cortical hierarchy ^29^. Then a series of analyses assessed spatial associations between functional connectivity distance and different features of neural organization, where 1,000 non-parametric spin tests ^99^ were performed to control spatial autocorrelation. We assessed relationships between functional connectivity distance and participation coefficient ^44^ and the functional gradient ^33^. To assess which cognitive faculties are related to long-and short-range connectivity, we analyzed meta-analytic z-statistics maps of 24 terms ^28,33^ from neurosynth (http://www.neurosynth.org) ^42^ covering a wide range of cognitive functions.

To examine associations to cortical microarchitecture, we assessed spatial associations between functional connectivity distance and *in vivo* qT1 measures in the same subjects, as well as *post mortem* histology and gene expression. We constructed 14 equivolumetric intracortical surfaces to sample qT1 intensities across cortical depths ^28,100,101^. Nodal microstructural profiles were cross-correlated across the cortical mantle to generate a microarchitectural similarity matrix and to derive the principal gradient of microstructural differentiation ^26,28,33^. In a separate analysis, we sampled intensities across cortical depths ^101^ of the BigBrain dataset, an ultra-high-resolution 3D volumetric histological reconstruction of a *post mortem* human brain ^50^. As in prior work ^102^, we estimated skewness of intensity profiles to represent microstructural laminar differentiation ^29,52^. In addition, we examined the transcriptomics data of the Allen Human Brain Atlas ^53^ to construct a gene co-expression matrix, and estimated the first component using principal component analysis following a recent analysis ^76^.

Finally, we assessed distance-based stratification of microarchitectural similarity based on *in vivo* imaging, histology, and gene co-expression data (**Fig. 3A**). A mixed effects model assessed distance effects on the combined qT1, BigBrain and gene co-expression features. In addition to studying effects globally, we also examined the association between microarchitectural similarity and functional connectivity distance across individual levels of the cortical hierarchy and in individual functional communities.

To verify the reproducibility of our findings, we repeated the main analyses on an independent validation dataset from the Human Connectome Project ^40^, including 200 unrelated healthy young adults (28.68 ± 3.69 years, 119 females).

## Acknowledgements

Yezhou Wang, Dr. Alan Evans, and Dr Bernhardt were supported by the Helmholtz International BigBrain Analytics and Learning Laboratory (HiBALL). Dr. Jessica Royer was supported by a Canadian Open Neuroscience Platform (CONP) fellowship and the Canadian Institute of Health Research (CIHR). Dr. Bo-yong Park was funded by the National Research Foundation of Korea (NRF-2021R1F1A1052303), Institute for Information and Communications Technology Planning and Evaluation (IITP) funded by the Korea Government (MSIT) (2020-0-01389, Artificial Intelligence Convergence Research Center, Inha University; 2021-0-02068, Artificial Intelligence Innovation Hub), and Institute for Basic Science (IBS-R015-D1). Sara Larivière was funded by CIHR and a Richard and Ann Sievers award. Reinder Vos de Wael was funded by studentships from the Savoy foundation for Epilepsy and a Richard and Ann Sievers award. Dr. Casey Paquola was funded through a postdoctoral fellowship of the FRQ-S. Dr. Sofie Valk was funded by the Max Planck Institute. Dr. Boris Bernhardt furthermore acknowledges research support from the National Science and Engineering Research Council of Canada (NSERC Discovery-1304413), the Canadian Institutes of Health Research (CIHR FDN-154298), SickKids Foundation (NI17-039), Azrieli Center for Autism Research (ACAR-TACC), BrainCanada (Future-Leaders), and the Tier-2 Canada Research Chairs (CRC) program. Data were provided, in part, by the Human Connectome Project, WU-Minn Consortium (Principal Investigators: David Van Essen and Kamil Ugurbil; 1U54MH091657) funded by the 16 NIH Institutes and Centers that support the NIH Blueprint for Neuroscience Research; and by the McDonnell Center for Systems Neuroscience at Washington University.

