## Supplementary Materials for "Long-range connections mirror and link microarchitectural and cognitive hierarchies in the human brain"

### SUPPLEMENTAL MATERIALS AND METHODS

#### Participants

We studied imaging and phenotypic data of 50 unrelated healthy adults (age: 29.82±5.73 years, 21 females) [1]. Data were collected between April 2018 and March 2020, and all participants denied a history of neurological illness. All participants had quality-controlled multimodal MRI data available (see below). The Ethics Committee of the Montreal Neurological Institute and Hospital approved the study. Written informed consent was obtained from all participants.

To verify the reproducibility of our findings, we repeated the main analyses on an independent validation dataset from the Human Connectome Project [2], including 200 unrelated healthy young adults (28.68 ± 3.69 years, 119 females).

#### MRI acquisition

MRI parameters are described in the MICA-MICs data descriptor [1]. In brief, scans were acquired using a 3T Siemens Magnetom Prisma-Fit equipped with a 64-channel head coil at the McConnell Brain Imaging Centre of the Montreal Neurological Institute. All participants underwent T1w structural MRI, multiband rs-fMRI, and qT1 imaging.

Two T1w scans with identical parameters were acquired with a 3D magnetization-prepared rapid gradient echo (MPRAGE) sequence (0.8mm isovoxels, matrix=320×320, 224 sagittal slices, repetition time (TR)=2300ms, echo time (TE)=3.14ms, inversion time (TI)s=900ms, flip angle=9°, iPAT=2). Scans were visually examined to ensure minimal head motion, and repeated if necessary. qT1 relaxometry data was acquired using a 3D-MP2RAGE sequence (0.8mm isovoxels, 240 sagittal slices, TR=5000ms, TE=2.9ms, TI 1=940ms, T1 2=2830ms, flip angle 1=4°, flip angle 2=5°, iPAT=3, bandwidth=270 Hz/px, echo spacing=7.2ms, partial Fourier=6/8). Two inversion images were combined for qT1 mapping to minimize sensitivity to B1 inhomogeneities and optimize intra- and inter-subject reliability[3,4].

A 7-minute rs-fMRI scan was acquired using multiband accelerated 2D-BOLD echo-planar imaging, EPI (TR=600ms, TE=30ms, 3mm isovoxels, flip angle=52°, FOV=240×240mm$^2$, slice thickness=3mm, mb factor=6, echo spacing=0.54ms). Participants were instructed to keep their eyes open, not fall asleep, and look at a fixation cross. Two spin-echo images with reverse phase encoding were also included for distortion correction of the rs-fMRI scans (phase encoding=AP/PA, 3mm isovoxels, FOV=240×240mm$^2$, slice thickness=3mm, TR=4029 ms, TE=48ms, flip angle=90°, echo spacing=0.54 ms, bandwidth= 2084 Hz/Px).

For HCP data, T1-weighted, T2-weighted, and rs-fMRI data were obtained using a Siemens Skyra 3T at Washington University [2]. The T1-weighted images were acquired using a MPRAGE sequence (TR=2,400 ms; TE=2.14 ms; FOV=224 × 224 mm$^2$; voxel size=0.7 mm$^3$; and number of slices=256). T2-weighted data were obtained with a T2-SPACE sequence, and acquisition parameters were aligned to those of the T1-weighted data except for TR (3,200 ms) and TE (565 ms). The rs-fMRI data were collected using a gradient-echo echo-planar imaging sequence (TR=720 ms; TE=33.1 ms; FOV=208 × 180 mm$^2$; voxel size=2 mm$^3$; number of slices=72; and number of volumes=1,200 per time series). During the rs-fMRI scan, participants were instructed to keep their eyes open looking at a fixation cross. Two sessions of rs-fMRI data were acquired; each of them contained data of left-to-right and right-to-left phase-encoded directions, providing up to four time series per participant.

##### Data preprocessing

Raw DICOMS were sorted using custom scripts, and converted to NIfTI using dcm2niix (https://github.com/rordenlab/dcm2niix)[5], renamed, and assigned to their respective subject-specific BIDS directories [6]. The BIDS validator (https://bids-standard.github.io/bids-validator/) ensured agreement to BIDS standards. Preprocessing was based on open-source scripts (http://github.com/MICA-MNI/micapipe).

T1w scans were deobliqued and reoriented, linearly co-registered, averaged, and corrected for intensity nonuniformity. Resulting volumes were skull stripped using FSL FIRST [7]. We used FreeSurfer 6.0[8,9] to generate cortical surface models from native T1w scans. Then, 14 equivolumetric surfaces were constructed for each participant between pial and white matter interfaces. These surfaces were used to systematically sample qT1 intensity profiles to construct individual microstructural profile similarity matrices.

The rs-fMRI data were pre-processed using AFNI[10] and FSL[7] tools. The first five volumes were discarded to ensure magnetic field saturation. Volumes were reoriented, followed by motion and distortion correction. Nuisance signal was removed using FMRIB's ICA-based X-noiseifier (ICA-FIX) [11] and by performing spike regression, which regressed out timepoints with large motion spikes [12,13]. Volume timeseries were registered to FreeSurfer space using boundary-based registration [14]. Surface-mapped timeseries were registered to Conte69, a template that has 32k vertices per hemisphere, and smoothed using a surface-based diffusion kernel (FWHM=10mm).

For HCP data, minimal preprocessing was performed using FSL, FreeSurfer, and Workbench [7,15,16]. T1- and T2-weighted data were corrected for gradient nonlinearity and b0 distortions, and then were co-registered using a rigid-body transformation. White and pial surfaces were generated using FreeSurfer [8,9,17]. A midthickness surface was generated by averaging white and pial surfaces, and used to generate the inflated surface that was registered to the Conte69 template [18] using MSMAll [19] and downsampled to a 32k vertex mesh. HCP provides a myelin-sensitive proxy based on the ratio of the T1- and T2-weighted contrast [20,21]. Then, rs-fMRI data were corrected for distortions and head motion, and were registered to the T1-weighted data and subsequently to MNI152 space. Magnetic field bias correction, skull removal, and intensity normalization were performed. Noise components were removed using ICA-FIX [11]. Time series were mapped to the standard grayordinate space, with a cortical ribbon-constrained volume-to-surface mapping algorithm.

##### Functional connectivity distance computation

Functional connectomes were generated by correlating preprocessed rs-fMRI averaged from 200 functionally defined cortical parcels [22]. Correlation matrices underwent Fisher-R-to-Z transformations. Individual geodesic distance matrices were calculated from the native pial surface using Surfdist (https://github.com/NeuroanatomyAndConnectivity/surfdist)[23]. Geodesic distance between cortical parcels was estimated by avoiding non-cortical paths (e.g. medial wall), and subsequently mapped to the native cortical surface and averaged within nodes. We averaged the within-hemisphere geodesic distance of both hemispheres to generate inter-hemispheric connections. For each region, we retained the top 10% of edges, and averaged the geodesic distance to all other regions in this configuration to estimate regional connectivity distance (**Fig. 1A**). We then calculated functional connectivity distance. For each region, we correlated its top 10% functional connections with the connectivity distance to assess distance-dependent functional network embedding. Here, positive values indicate increased embedding of a region with long-range connectivity networks, while negative values indicate embedding with short range connectivity networks. Subject-level data were averaged to generate group-level estimates. In addition to surface mapping, we stratified functional connectivity distance according to intrinsic functional communities [24] and a model of primate cortical hierarchy [25] (**Fig. 1B, *SI Appendix*, Fig. S1**). To verify the robustness of functional connectivity distance construction, we constructed it using different thresholds of functional connectivity (5%, 15%, 20%, 25%), as well as functional data with GSR (***SI Appendix*, Fig. S2**). We furthermore evaluated inter-individual variations of functional connectivity distance and functional associations (***SI Appendix*, Fig. S3**).

##### Functional and microarchitectural contextualization

A series of analyses assessed spatial associations between functional connectivity distance and different features of neural organization. Unless otherwise specified, 1,000 non-parametric spin tests [26] assessed spatial associations while controlling for spatial autocorrelation.

*Functional contextualization.* We first assessed the spatial correlation between functional connectivity distance and the functional participation coefficient[27], a graph theoretical index of connectional diversity. [28] (***SI Appendix*, Fig. S1**). The participation coefficient was computed at both group- and individual-level, where each brain region was assigned to one of seven predefined functional communities [24]. For a given region, its participation coefficient will be 0 if its connections are entirely restricted to one given community, which estimated by intrinsic functional connectivity [24]. In contrast, participation coefficient will approach 1 if the region's connections are evenly distributed among all communities. We also assessed associations between functional connectivity distance and the principal functional gradient [28]. Functional gradients were calculated using BrainSpace (https://github.com/MICA-MNI/BrainSpace) [29]. In brief, an individual's functional connectome was thresholded to retain the top 10% connectivity per region, and this connectome was converted into a normalized angle matrix. The principal functional gradient was identified using diffusion map embedding [30], an efficient nonlinear dimensionality reduction technique that is robust to noise [31,32]. It is controlled by two parameters $\alpha$ and $t$, where $\alpha$ controls influence of the density of sampling points on the manifold ($\alpha=1$, no influence; $\alpha=0$, maximal influence), and $t$ controls the scale of eigenvalues of the diffusion operator. We set $\alpha = 0.5$ and $t = 0$ based on recommendations to retain the global relations between data points in the embedded space[28,29,33,34]. Individual-level manifolds were aligned to a template manifold obtained from the HCP dataset via Procrustes alignment[35], and averaged to generate a group-level manifold (**Fig. 1B**).

Meta-analysis on cognitive functions. To assess which cognitive faculties are related to long- and short-range connectivity, we analyzed meta-analytic maps from Neurosynth (http://www.neurosynth.org) [36]. Neurosynth combines meta-analysis and text mining techniques to generate probabilistic mappings between cognitive terms and spatial brain patterns. We analyzed meta-analytic z-statistics maps of 24 terms [28,33] covering a wide range of cognitive functions (**Fig. 1C**). The vertex-level z-statistic was mapped to the Schaefer atlas [22] with 200 regions, and averaged within each region. We divided the functional connectivity distance into ten-percentile bins and calculated the mean z-statistic of per term within each bin. This resulted in overall z-activations of each cognitive term with respect to functional connectivity distance. Findings were replicated on the HCP data set (***SI Appendix*, Fig. S4**).

Microarchitectural contextualization. We assessed spatial associations to *in vivo* qT1 measures in the same subjects, as well as *post mortem* histology and gene expression (**Fig. 2**).

qT1. Consistent with previous work[33,37,38], we constructed 14 equivolumetric intracortical surfaces to sample qT1 intensities across cortical depths. Data sampled from surfaces closest to the pial and white matter boundaries were removed to mitigate partial volume effects. The fsaverage5 surface-mapped Schaefer atlas with 200 parcels [22] was interpolated to the native surface in each individual, and vertex-wise intensity profiles were averaged within parcels. Nodal microstructural profiles were cross-correlated across the cortical mantle using partial correlations, while controlling for the average cortex-wide intensity profile. The resulting matrix was thresholded and log-transformed. This matrix captures the similarity in intracortical microstructural profiles across the cortex. In line with prior work and the functional analyses [28,33,39], we thresholded the microstructural similarity matrix, converted it into a normalized angle matrix, and derived microarchitectural gradients using diffusion map embedding. Individual-level gradients were aligned to the template manifold obtained from the group-level microstructural similarity matrix, and averaged to generate a group-level microstructural gradient[29]. The association between functional connectivity distance and the principal microarchitectural gradient derived from qT1 measures was examined both at the group and individual level (**Fig. 2A; *SI Appendix*, Fig. S5A**). Analyses were repeated in the HCP dataset (***SI Appendix*, Fig. S6**), which uses T1w/T2w imaging ratio as an alternative microstructural index. We also assessed skewness of microstructural profiles across cortical layers and averaged them across all participants to contrast relative properties of deep and superficial cortical layers. Intracortical depth is a critical dimension of laminar differentiation that relates to architectural complexity [40] and cortical hierarchy[25]. Correlations between functional connectivity distance and qT1 skewness were examined on individual-level and group-level (***SI Appendix*, Fig. S5B**).

BigBrain. BigBrain is an ultra-high-resolution 3D volumetric histological reconstruction of a *post mortem* human brain (https://bigbrain-ftp.loris.ca/) [41]. We constructed 16 equivolumetric intracortical surfaces with 163,842 matched vertices per hemisphere to sample intensities across cortical depths[38], yielding intensity profiles reflecting microstructural composition of each cortical vertex. Data sampled from surfaces closest to the pial and white matter boundaries were discarded to mitigate partial volume effects. Using SurfStat for Matlab (http://mica-mni.github.io/surfstat), we constructed surface-based linear models and controlled the midsurface *y* coordinate to account for an anterior-posterior increase in intensity values across the BigBrain due to coronal slicing and reconstruction[33]. Vertex-wise intensity profiles were subsequently averaged within functionally defined 200 parcels [22]. We then estimated skewness of intensity profiles to represent microstructural differentiation [25,40], and assessed the association to functional connectivity distance (**Fig. 2B**).

Gene expression. We examined microarray gene expression data provided by the Allen Human Brain Atlas (AHBA) [43]. Among all genes from AHBA, we downsampled the expression data to a nodal level and then selected only genes which were consistently expressed across 6 donors using abagen (https://github.com/rmarkello/abagen)[44]. For 4 donors who lacked expression data of right hemisphere, used their left hemisphere to replace. For each gene, the whole-brain gene expression maps between all pairs of donors were correlated, while genes with an average inter-donor $r \leq 0.5$ were discarded (**Fig. 2C**). We then constructed a gene co-expression matrix by cross correlating these genes, and estimated the first component of gene co-expression using principal component analysis, a linear dimensionality reduction technique. Next, we correlated the first principal component to the functional connectivity distance map, controlling for spatial autocorrelation. These selected genes were then spatially correlated with the functional connectivity distance map (false discovery rate (FDR) correction [45]), to further investigated genes whose expression correlated to functional connectivity distance (FDR $< 0.05$; ***SI Appendix***, **Table S2**). We only focused on genes that were positively correlated to longer functional connectivity distance. To investigate cell-type-specific gene enrichment, we compared these genes to cell-type specific genes, including excitatory and inhibitory neurons as well as microglia, endothelial cells, pericytes, astrocytes, as well as oligodendrocytes and precursor cells [46,47]. To assess the distribution of genes in each cell type, we calculated the overlap ratio (***SI Appendix***, **Fig. S7**). Here, spin tests and FDR were used again to correct findings for multiple comparisons and spatial autocorrelation.

**Distance based stratification of microarchitectural similarity**

For each cortical parcel, we retained the top 10% of edges in the group averaged functional connectivity matrix and averaged the microarchitectural similarity of this network (based on qT1, BigBrain, gene co-expression; **Fig. 3A**). Parcels were then sorted relative to functional connectivity distance, and we fitted linear as well as quadratic functions to model this relationship (***SI Appendix***, **Fig. S8**). In addition to assessing distance effects on a single metric, we also built a mixed effects model that assesses distance effects on the microarchitectural similarity based on qT1, BigBrain, and gene expression. Analyses were carried out across the whole cortex, and repeated in specific functional communities and cortical hierarchy levels (**Fig. 3B**, ***Appendix***, **Fig. S9**). We repeated the main analyses on HCP dataset and found consistent findings (***SI Appendix***, **Fig. S10**).

#### SUPPLEMENTARY FIGURES

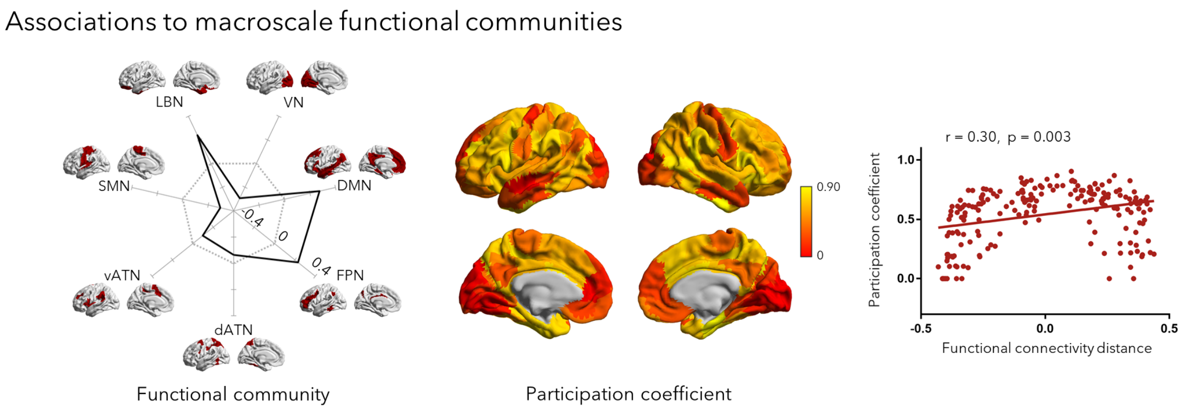


**Fig. S1. Associations to macroscale functional communities.** Stratification of functional connectivity distance across macroscale functional communities [41], and associations to the functional participation coefficient.

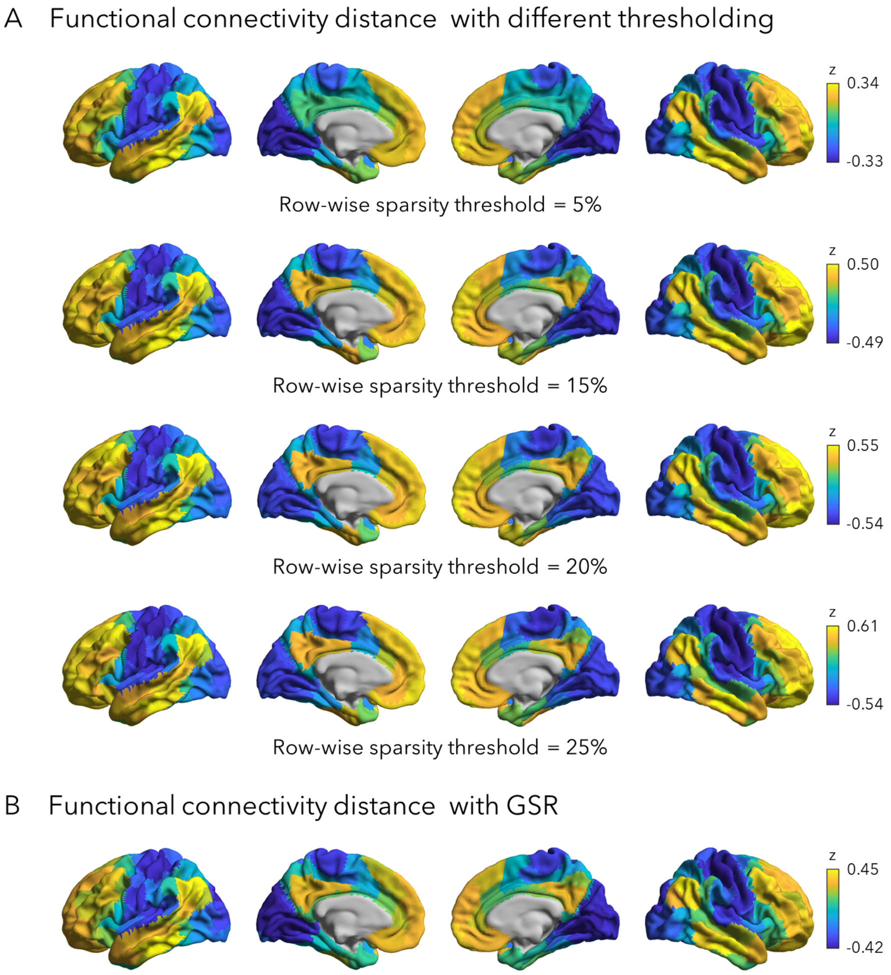


**Fig. S2. Robustness analysis on functional connectivity distance. (A)** Functional connectivity distance computation across different functional connectome thresholdings. **(B)** Functional connectivity distance after additional global signal regression (GSR).

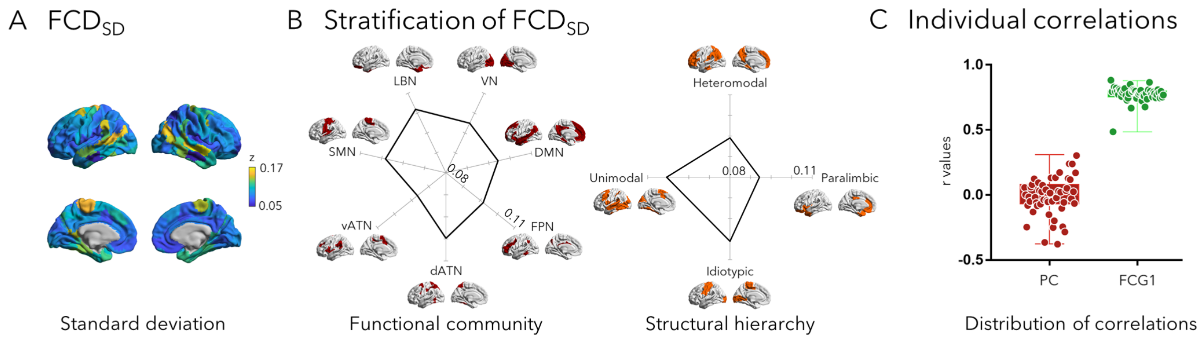


**Fig. S3. Individual-level functional connectivity distance characteristics. (A)** Standard deviation of functional connectivity distance ($FCD_{SD}$) across all individuals. **(B)** Stratification of $FCD_{SD}$ on functional communities and structural hierarchies. **(C)** Distribution of correlations between functional connectivity distance and other features (participation coefficient and principal gradient of functional connectome) on individual level. *Abbreviation*: PC, participation coefficient; FCG1, principal gradient of functional connectome.

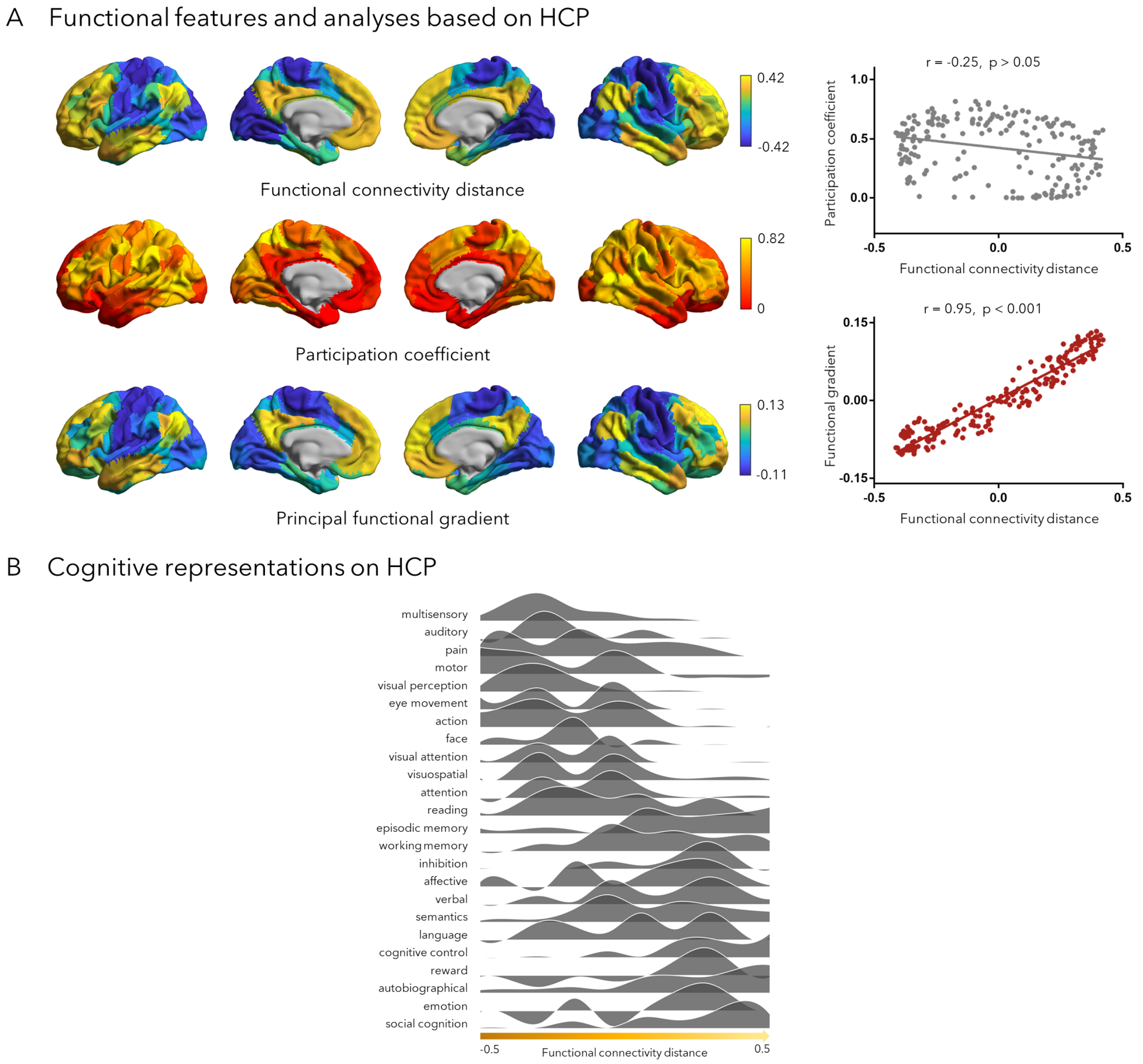


**Fig. S4. Replication analyses based on HCP. (A)** Functional features and analyses based on the HCP dataset. *Left panel*: Group-level functional connectivity distance, participation coefficient, and principal functional gradient. *Right panel:* Associations between participation coefficient, principal functional gradient, and functional connectivity distance. **(B)** Cognitive associations. Distributions of meta-analytical task-related fMRI activation across cortical regions relative to functional connectivity distance.

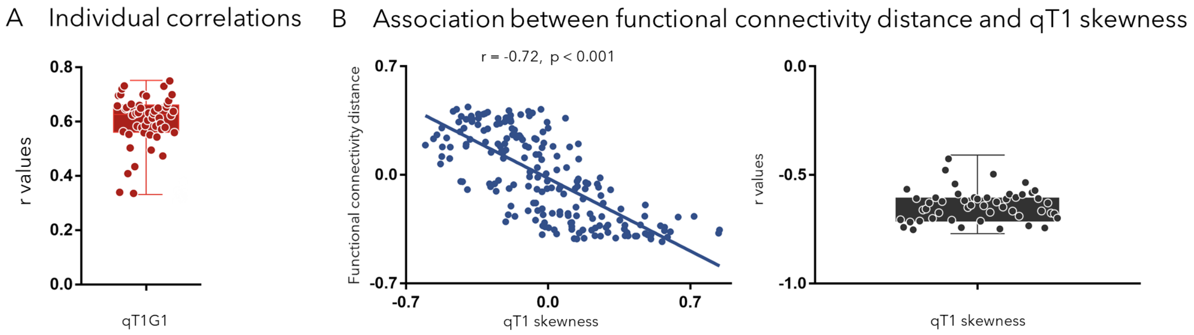


**Fig. S5. Correlations between functional connectivity distance and in vivo microstructural features. (A)** Individual-level correlations between functional connectivity distance and microstructural gradients derived from qT1. **(B)** Association between functional connectivity distance and microstructure skewness derived from qT1 at a group and individual level. *Abbreviation*: qT1G1, microstructure gradient derived from qT1.

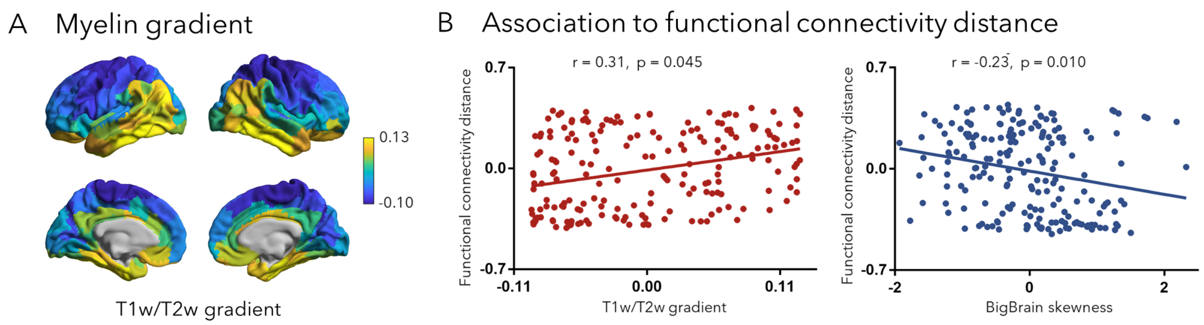


**Fig. S6. Structure-function association in the HCP dataset. (A)**: Principal microstructural gradient estimated based on T1w/T2w ratios. **(B)** Association between functional connectivity distance and microstructure gradients derived from T1w/T2w, and BigBrain skewness feature.

C Cell-type specific expression analysis

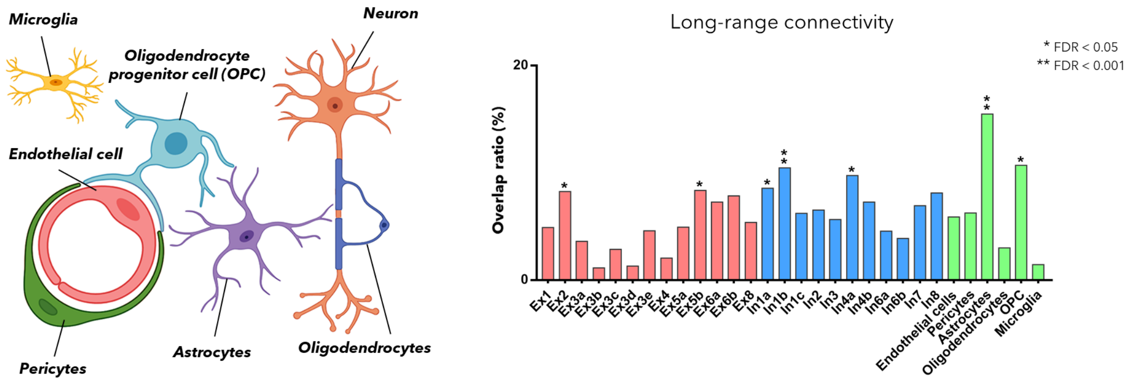


**Fig. S7. Cell-type specific gene expression analysis.** *Left*: Schema of different cell types. *Right*: The overlap ratio between genes positively related to functional connectivity distance (long-range connectivity) and cell-type specific genes. Supragranular neurons: Ex1, Ex2, Ex3a, Ex3b, Ex3c, Ex3d, Ex3e, In1a, In1b, In1c, In2, In3; infragranular neurons: Ex4, Ex5a, Ex5b, Ex6a, Ex6b, Ex8, In4a, In4b, In6a, In6b, In7, In8. *Abbreviation*: FDR, false discovery rate; Ex, excitatory neuron; In, inhibitory neuron.

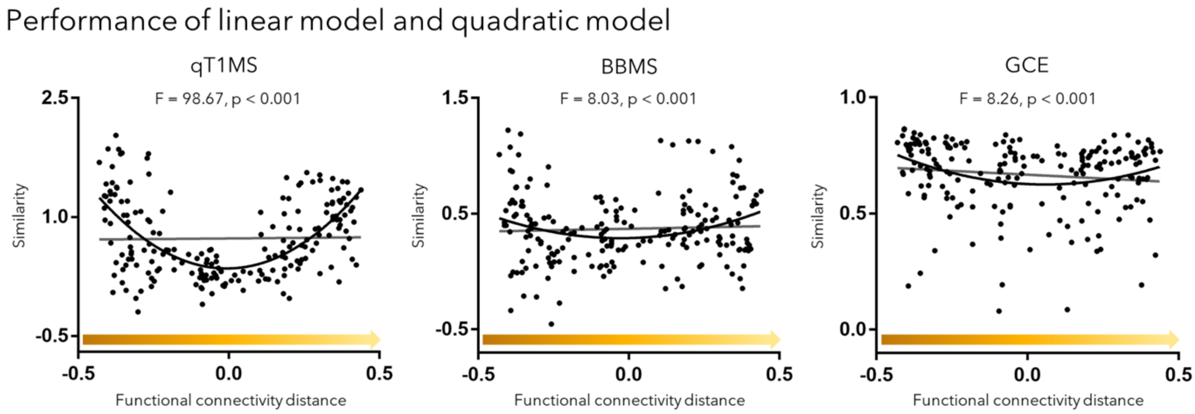


**Fig. S8. Performance of linear model and quadratic model.** Linear model (*gray line*) and quadratic model (*black curve*) were used to fit the association between microarchitectural similarity and functional connectivity distance. The golden arrows whose color change from darker to lighter represent the gradual increase in functional connectivity distance.

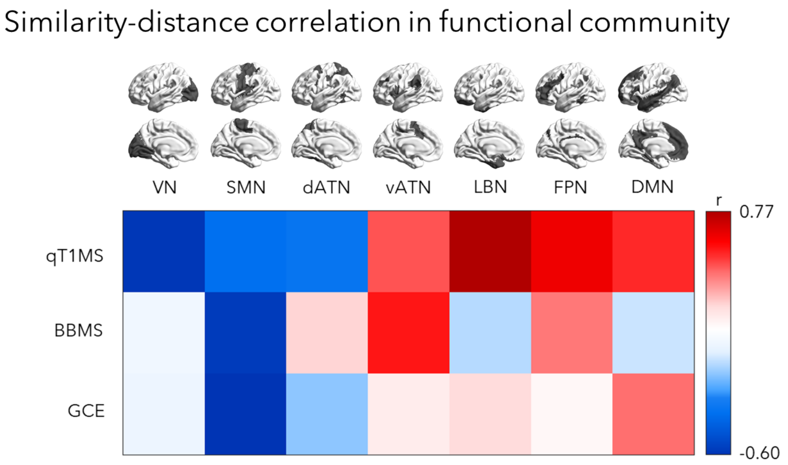


**Fig. S9. Similarity-distance correlation in functional community.** The correlation between microarchitectural similarity and functional connectivity distance was examined across different intrinsic functional communities. Different colors in the matrix represent correlations between distance and microstructural/genetic similarities (r values). *Abbreviation*: VN, visual network; SMN, somatomotor network; dATN, dorsal attention network; vATN, ventral attention network; LBN, limbic network; FPN, frontoparietal control network; DMN, default mode network.

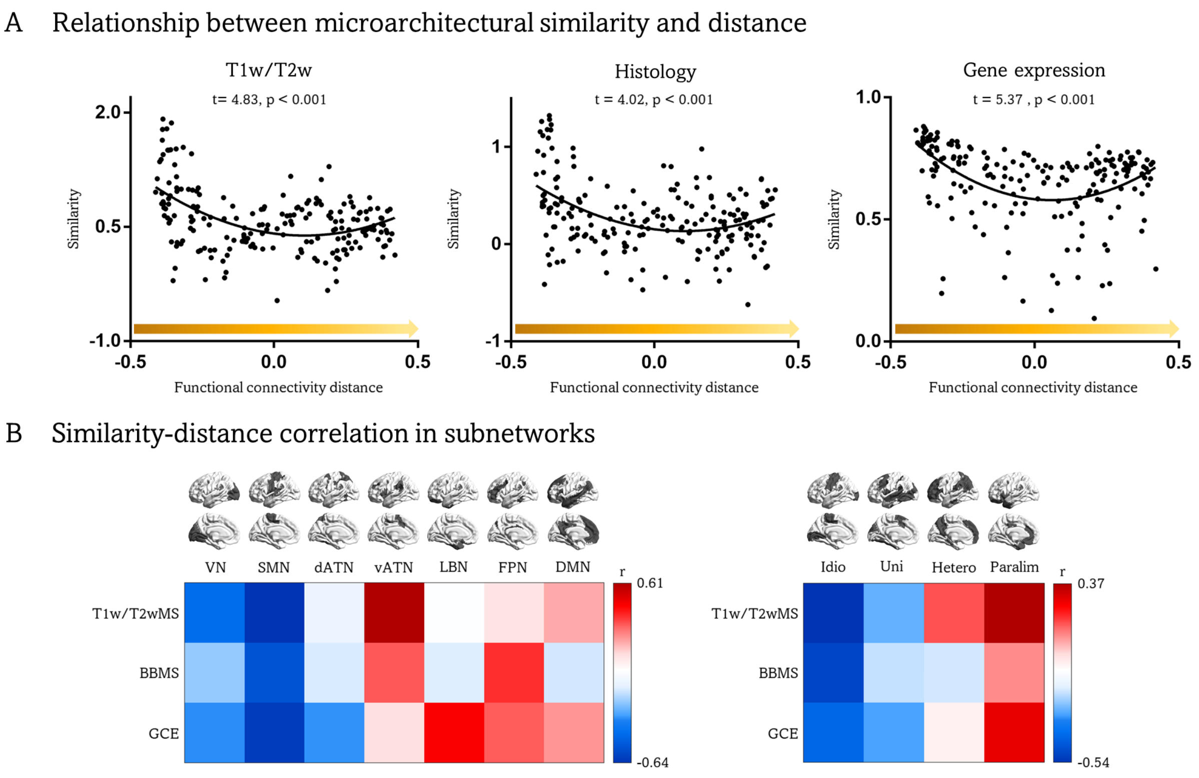


**Fig. S10. Microarchitectural and genetic similarity analyses based on the HCP dataset. (A)** Quadratic relationship between microarchitectural similarity and functional connectivity distance. (**B)** Similarity-distance associations in intrinsic functional communities and levels of the cortical hierarchy. Different colors in the matrix represent correlations between functional connectivity distance and microarchitectural similarity (r values). *Abbreviation*: T1w/T2wMS, microstructural similarity derived from T1w/T2w; BBMS, microstructural similarity derived from BigBrain; GCE, gene co-expression; VN, visual network; SMN, somatomotor network; dATN, dorsal attention network; vATN, ventral attention network; LBN, limbic network; FPN, frontoparietal control network; DMN, default mode network.
