## Supplementary material for "Long-range connections mirror and link microarchitectural and cognitive hierarchies in the human brain": Table S1

**Table S1.** Original gene list with genes co-expressed across all donors

| Gene | r Value |
|---|---|
| ABCA6 | 0.55 |
| ABTB1 | 0.50 |
| ACAN | 0.74 |
| ACSL6 | 0.52 |
| ACTC1 | 0.53 |
| ACTG1 | 0.54 |
| ACTR1B | 0.63 |
| ACVR2A | 0.55 |
| ACYP2 | 0.55 |
| ADAM22 | 0.54 |
| ADAM23 | 0.58 |
| ADAMTS19 | 0.52 |
| ADAMTS3 | 0.71 |
| ADCY7 | 0.58 |
| ADCYAP1R1 | 0.57 |
| ADGRG1 | 0.70 |
| ADGRL3 | 0.56 |
| ADGRV1 | 0.59 |
| ADM | 0.56 |
| ADO | 0.66 |
| ADPRHL1 | 0.52 |
| ADRA1B | 0.51 |
| ADTRP | 0.57 |
| AFTPH | 0.60 |
| AHI1 | 0.64 |
| AHRR | 0.53 |
| AK4 | 0.73 |
| AKAIN1 | 0.59 |
| AKAP14 | 0.54 |
| AKAP7 | 0.56 |
| AKR1C3 | 0.50 |
| ALDH1A3 | 0.76 |
| ALDH4A1 | 0.53 |
| ALDH6A1 | 0.51 |
| ALKAL2 | 0.65 |
| AMDHD1 | 0.54 |
| AMIGO2 | 0.70 |
| ANK1 | 0.75 |
| ANKDD1A | 0.51 |
| ANKH | 0.58 |
| ANKRD19P | 0.62 |
| ANKRD29 | 0.52 |
| ANKRD34C | 0.58 |
| ANKRD50 | 0.70 |
| ANKRD6 | 0.71 |
| ANKS6 | 0.52 |
| ANO3 | 0.59 |
| AP1S2 | 0.50 |

| | |
|---|---|
| AP2B1 | 0.50 |
| APCDD1 | 0.51 |
| APOC1 | 0.59 |
| APOE | 0.54 |
| AQP4 | 0.53 |
| AQP9 | 0.59 |
| AR | 0.69 |
| ARHGAP12 | 0.59 |
| ARHGAP25 | 0.58 |
| ARHGAP28 | 0.64 |
| ARHGAP6 | 0.59 |
| ARHGAP9 | 0.65 |
| ARHGDIA | 0.55 |
| ARHGEF28 | 0.53 |
| ARID2 | 0.54 |
| ARID3B | 0.54 |
| ARL15 | 0.54 |
| ARL9 | 0.55 |
| ARPP21 | 0.51 |
| ARRB1 | 0.53 |
| ASB13 | 0.76 |
| ASB2 | 0.56 |
| ASCL2 | 0.65 |
| ASGR2 | 0.51 |
| ASTN1 | 0.55 |
| ASXL3 | 0.50 |
| ATG4D | 0.57 |
| ATOH7 | 0.66 |
| ATP13A4 | 0.51 |
| ATP1A2 | 0.55 |
| ATP1B2 | 0.54 |
| ATP2B4 | 0.76 |
| AVPI1 | 0.58 |
| B4GALT2 | 0.55 |
| B9D1 | 0.58 |
| BAIAP2L2 | 0.51 |
| BAIAP3 | 0.78 |
| BBOX1 | 0.53 |
| BCAS1 | 0.50 |
| BCL11B | 0.52 |
| BEND6 | 0.58 |
| BHLHE22 | 0.69 |
| BHMT2 | 0.54 |
| BICDL2 | 0.59 |
| BID | 0.62 |
| BIRC3 | 0.70 |
| BLMH | 0.59 |
| BMP4 | 0.62 |
| BRINP1 | 0.61 |
| BTBD17 | 0.51 |
| C11orf97 | 0.58 |
| C14orf132 | 0.52 |

| | |
|---|---|
| C17orf75 | 0.59 |
| C1QL3 | 0.57 |
| C1R | 0.67 |
| C1S | 0.70 |
| C1orf61 | 0.54 |
| C2CD4C | 0.75 |
| C3orf18 | 0.53 |
| C5orf30 | 0.58 |
| C5orf49 | 0.51 |
| C6orf106 | 0.60 |
| C9orf3 | 0.58 |
| CABP1 | 0.59 |
| CACNA1H | 0.64 |
| CACNA2D2 | 0.53 |
| CACNB4 | 0.60 |
| CADM1 | 0.72 |
| CADPS2 | 0.73 |
| CALN1 | 0.55 |
| CAMK2D | 0.66 |
| CAMK2G | 0.51 |
| CARTPT | 0.62 |
| CASC4 | 0.51 |
| CASK | 0.50 |
| CASTOR1 | 0.56 |
| CBLN2 | 0.53 |
| CCBE1 | 0.68 |
| CCDC151 | 0.51 |
| CCDC25 | 0.51 |
| CCDC28B | 0.56 |
| CCDC39 | 0.61 |
| CCDC58 | 0.51 |
| CCDC8 | 0.51 |
| CCDC85C | 0.51 |
| CCDC90B | 0.61 |
| CCNI | 0.71 |
| CD24 | 0.72 |
| CD274 | 0.50 |
| CD6 | 0.57 |
| CD63 | 0.62 |
| CDC42EP4 | 0.50 |
| CDH11 | 0.56 |
| CDH13 | 0.53 |
| CDH7 | 0.52 |
| CDR2L | 0.60 |
| CDS1 | 0.74 |
| CEND1 | 0.59 |
| CENPJ | 0.66 |
| CENPVL3 | 0.60 |
| CENPW | 0.53 |
| CEP57 | 0.54 |
| CERK | 0.59 |
| CFAP44 | 0.58 |

| | |
|---|---|
| CFAP58-AS1 | 0.53 |
| CFD | 0.61 |
| CHAF1A | 0.66 |
| CHCHD6 | 0.58 |
| CHGA | 0.57 |
| CHID1 | 0.55 |
| CHMP1A | 0.56 |
| CHRDL1 | 0.60 |
| CHRNA2 | 0.51 |
| CHST15 | 0.57 |
| CHST7 | 0.55 |
| CHST9 | 0.53 |
| CIDEA | 0.58 |
| CITED2 | 0.62 |
| CKMT1B | 0.51 |
| CLDN10 | 0.54 |
| CLEC2L | 0.52 |
| CLEC4G | 0.51 |
| CLMP | 0.67 |
| CMAHP | 0.58 |
| CMYA5 | 0.53 |
| CNN3 | 0.63 |
| CNR1 | 0.65 |
| CNTN1 | 0.60 |
| CNTN3 | 0.52 |
| CNTN6 | 0.59 |
| COCH | 0.60 |
| COL11A1 | 0.53 |
| COL5A1 | 0.64 |
| COL5A2 | 0.55 |
| COPA | 0.52 |
| COX7A1 | 0.68 |
| CPE | 0.59 |
| CPLX1 | 0.66 |
| CPNE3 | 0.59 |
| CPNE6 | 0.69 |
| CPNE7 | 0.61 |
| CPNE9 | 0.64 |
| CPT1A | 0.55 |
| CRB1 | 0.52 |
| CREM | 0.58 |
| CRIM1 | 0.58 |
| CRTAC1 | 0.60 |
| CTNND2 | 0.63 |
| CTSK | 0.53 |
| CTSO | 0.55 |
| CTXN1 | 0.56 |
| CTXN3 | 0.71 |
| CUX1 | 0.56 |
| CXorf57 | 0.72 |
| CYP1B1 | 0.59 |
| DACH1 | 0.54 |

| | |
|---|---|
| DACH2 | 0.57 |
| DAPL1 | 0.54 |
| DCAF10 | 0.50 |
| DCBLD2 | 0.68 |
| DCN | 0.54 |
| DCP1A | 0.64 |
| DCUN1D2 | 0.74 |
| DDA1 | 0.65 |
| DDAH1 | 0.55 |
| DDAH2 | 0.61 |
| DDHD2 | 0.61 |
| DEFB131A | 0.53 |
| DENND1B | 0.57 |
| DENND2A | 0.52 |
| DENND2D | 0.58 |
| DERL1 | 0.51 |
| DEXI | 0.69 |
| DHDH | 0.60 |
| DIAPH2 | 0.53 |
| DIAPH3 | 0.63 |
| DIRAS3 | 0.59 |
| DIS3L2 | 0.59 |
| DKFZp779M0652 | 0.68 |
| DLC1 | 0.58 |
| DMKN | 0.60 |
| DNAAF2 | 0.52 |
| DNAAF4 | 0.51 |
| DNAH14 | 0.69 |
| DNAH5 | 0.51 |
| DNAJA4 | 0.67 |
| DNAJC21 | 0.60 |
| DNAJC4 | 0.61 |
| DOC2B | 0.59 |
| DOCK7 | 0.60 |
| DOK6 | 0.52 |
| DPP8 | 0.53 |
| DPY19L2P1 | 0.62 |
| DPY19L2P4 | 0.51 |
| DPYSL3 | 0.73 |
| DSCC1 | 0.56 |
| DTWD1 | 0.51 |
| DUS1L | 0.56 |
| DUSP1 | 0.50 |
| DYDC2 | 0.65 |
| DYNLL1 | 0.57 |
| E2F5 | 0.55 |
| ECSIT | 0.57 |
| EDNRB | 0.63 |
| EFCAB1 | 0.78 |
| EFEMP1 | 0.53 |
| EFHC2 | 0.61 |
| EFNA5 | 0.56 |

| | |
|---|---|
| EGLN1 | 0.57 |
| EIF2A | 0.52 |
| EIF4E1B | 0.66 |
| EIF5A2 | 0.64 |
| ELMO3 | 0.65 |
| ELOVL4 | 0.50 |
| EMILIN2 | 0.57 |
| EML2 | 0.50 |
| ENKUR | 0.56 |
| ENO2 | 0.51 |
| ENOX1 | 0.70 |
| ENTPD4 | 0.61 |
| EPB41 | 0.63 |
| EPB41L4B | 0.53 |
| EPC2 | 0.56 |
| EPCAM | 0.60 |
| EPHX1 | 0.55 |
| EPN3 | 0.78 |
| ERICH5 | 0.55 |
| ERICH6-AS1 | 0.63 |
| ERLIN2 | 0.51 |
| ERRFI1 | 0.60 |
| ESRRA | 0.69 |
| ESRRG | 0.81 |
| ESYT1 | 0.52 |
| ETNPPL | 0.63 |
| EXOC6 | 0.55 |
| EXTL2 | 0.54 |
| EYA2 | 0.57 |
| F12 | 0.56 |
| F3 | 0.53 |
| FABP5P3 | 0.59 |
| FABP7 | 0.61 |
| FAM107A | 0.56 |
| FAM110A | 0.74 |
| FAM110C | 0.61 |
| FAM120AOS | 0.53 |
| FAM122C | 0.51 |
| FAM131B | 0.56 |
| FAM135B | 0.55 |
| FAM149A | 0.60 |
| FAM171B | 0.73 |
| FAM181A | 0.64 |
| FAM189B | 0.51 |
| FAM196A | 0.54 |
| FAM20A | 0.56 |
| FAM210B | 0.52 |
| FAM213A | 0.59 |
| FAM214A | 0.55 |
| FAM216A | 0.73 |
| FAM234B | 0.55 |
| FAM43A | 0.58 |

| | |
|---|---|
| FAM49B | 0.61 |
| FAM57B | 0.67 |
| FAM71E1 | 0.71 |
| FAM71F1 | 0.63 |
| FAM78B | 0.53 |
| FANCI | 0.55 |
| FARP1 | 0.58 |
| FASTKD5 | 0.56 |
| FBLN2 | 0.51 |
| FBLN7 | 0.71 |
| FBXO33 | 0.64 |
| FBXO9 | 0.58 |
| FER1L4 | 0.52 |
| FERMT2 | 0.65 |
| FES | 0.67 |
| FGF18 | 0.59 |
| FGF2 | 0.64 |
| FGF9 | 0.80 |
| FGFR3 | 0.54 |
| FIBIN | 0.64 |
| FLRT3 | 0.54 |
| FLT3 | 0.58 |
| FNBP1L | 0.72 |
| FNDC4 | 0.62 |
| FNDC5 | 0.69 |
| FRAT1 | 0.57 |
| FRAT2 | 0.53 |
| FREM3 | 0.62 |
| FSTL1 | 0.60 |
| FSTL5 | 0.53 |
| FUOM | 0.51 |
| FXYD1 | 0.52 |
| FZD1 | 0.61 |
| FZD8 | 0.56 |
| GABRA1 | 0.60 |
| GABRA2 | 0.55 |
| GABRA3 | 0.62 |
| GABRA5 | 0.79 |
| GABRB1 | 0.63 |
| GABRD | 0.60 |
| GABRE | 0.54 |
| GABRG1 | 0.63 |
| GAL | 0.64 |
| GALNTL5 | 0.51 |
| GAP43 | 0.53 |
| GASAL1 | 0.51 |
| GDF10 | 0.68 |
| GDPD1 | 0.58 |
| GGCT | 0.56 |
| GGTA1P | 0.58 |
| GIGYF1 | 0.50 |
| GJA1 | 0.59 |

| | |
|---|---|
| GJB6 | 0.53 |
| GLCCI1 | 0.78 |
| GLOD4 | 0.61 |
| GLRA3 | 0.68 |
| GLRX | 0.55 |
| GLS2 | 0.67 |
| GLUD1 | 0.58 |
| GLUD2 | 0.60 |
| GMFB | 0.70 |
| GMPR | 0.53 |
| GNAQ | 0.56 |
| GNB4 | 0.71 |
| GNG10 | 0.59 |
| GNG12 | 0.55 |
| GNG2 | 0.71 |
| GNG4 | 0.60 |
| GNPTAB | 0.54 |
| GOLM1 | 0.59 |
| GPAT3 | 0.83 |
| GPC4 | 0.62 |
| GPC5 | 0.60 |
| GPCPD1 | 0.75 |
| GPD2 | 0.63 |
| GPHN | 0.56 |
| GPLD1 | 0.70 |
| GPR137C | 0.56 |
| GPR143 | 0.53 |
| GPR160 | 0.73 |
| GPR161 | 0.63 |
| GPR68 | 0.66 |
| GPR88 | 0.54 |
| GPX3 | 0.70 |
| GRIA1 | 0.70 |
| GRIK2 | 0.57 |
| GRP | 0.65 |
| GSDMB | 0.57 |
| GSG1 | 0.50 |
| GSTM2 | 0.53 |
| GSTM5 | 0.50 |
| GSTT1 | 0.68 |
| GUCA2B | 0.63 |
| GUCY1A2 | 0.55 |
| HAPLN4 | 0.73 |
| HCG23 | 0.50 |
| HCG4 | 0.56 |
| HCN1 | 0.50 |
| HDC | 0.54 |
| HEATR3 | 0.50 |
| HERPUD1 | 0.50 |
| HES5 | 0.61 |
| HHIPL1 | 0.53 |
| HIST1H1D | 0.63 |

| | |
|---|---|
| HIST1H4A | 0.57 |
| HIVEP2 | 0.62 |
| HMGCS1 | 0.51 |
| HPCAL4 | 0.68 |
| HR | 0.64 |
| HRASLS | 0.55 |
| HS3ST1 | 0.72 |
| HSPA4L | 0.60 |
| HSPB8 | 0.60 |
| HTR1A | 0.55 |
| HTR2C | 0.68 |
| HYLS1 | 0.53 |
| ICA1 | 0.54 |
| IDI2-AS1 | 0.53 |
| IER2 | 0.66 |
| IFFO1 | 0.58 |
| IFFO2 | 0.60 |
| IFNLR1 | 0.68 |
| IGDCC3 | 0.59 |
| IGFBP2 | 0.54 |
| IGFBP5 | 0.52 |
| IGIP | 0.52 |
| IL13RA2 | 0.72 |
| IL17RD | 0.54 |
| IL33 | 0.53 |
| IMPACT | 0.53 |
| INA | 0.61 |
| INIP | 0.56 |
| INTS8 | 0.60 |
| INTU | 0.59 |
| IPW | 0.57 |
| IQCA1 | 0.67 |
| IQCJ | 0.55 |
| IRS1 | 0.57 |
| ISG15 | 0.64 |
| ISOC1 | 0.57 |
| ITGA11 | 0.57 |
| ITGB1BP1 | 0.51 |
| ITPR1 | 0.55 |
| ITPRIPL2 | 0.60 |
| JADE1 | 0.55 |
| JAK2 | 0.52 |
| JDP2 | 0.61 |
| KANK4 | 0.52 |
| KBTBD2 | 0.56 |
| KBTBD6 | 0.53 |
| KCNA1 | 0.79 |
| KCNA2 | 0.71 |
| KCNAB3 | 0.76 |
| KCNC1 | 0.75 |
| KCNC3 | 0.65 |
| KCNG1 | 0.69 |

| | |
|---|---|
| KCNG3 | 0.57 |
| KCNH7 | 0.53 |
| KCNJ11 | 0.54 |
| KCNMB4 | 0.63 |
| KCNN3 | 0.70 |
| KCNS1 | 0.50 |
| KCNT1 | 0.56 |
| KCTD15 | 0.55 |
| KCTD4 | 0.75 |
| KCTD9 | 0.64 |
| KDF1 | 0.70 |
| KDM7A | 0.64 |
| KIAA1107 | 0.59 |
| KIF15 | 0.57 |
| KIF2C | 0.51 |
| KIRREL3 | 0.53 |
| KLF12 | 0.51 |
| KLF9 | 0.65 |
| KLHL13 | 0.53 |
| KLHL8 | 0.60 |
| KLK7 | 0.52 |
| KLRC2 | 0.54 |
| KNG1 | 0.59 |
| KRBOX4 | 0.59 |
| KRT31 | 0.52 |
| L2HGDH | 0.55 |
| L3MBTL4 | 0.52 |
| LAG3 | 0.82 |
| LAMA1 | 0.57 |
| LAMA2 | 0.60 |
| LAMA4 | 0.51 |
| LAPTM4B | 0.55 |
| LCA5 | 0.56 |
| LCP2 | 0.66 |
| LDHD | 0.53 |
| LEAP2 | 0.51 |
| LEPROTL1 | 0.54 |
| LGALS1 | 0.63 |
| LGI4 | 0.53 |
| LINC00260 | 0.63 |
| LINC00473 | 0.64 |
| LINC00515 | 0.58 |
| LINC00599 | 0.66 |
| LINC01137 | 0.59 |
| LINC01140 | 0.71 |
| LINC01296 | 0.69 |
| LINC01637 | 0.66 |
| LINC02223 | 0.54 |
| LIX1 | 0.60 |
| LMO1 | 0.61 |
| LMO3 | 0.50 |
| LOC100129291 | 0.59 |

| | |
|---|---|
| LOC100506124 | 0.57 |
| LOC285097 | 0.61 |
| LOC440934 | 0.53 |
| LOC642852 | 0.66 |
| LONRF3 | 0.53 |
| LOXL1 | 0.64 |
| LRCH1 | 0.56 |
| LRP1B | 0.60 |
| LRP4 | 0.55 |
| LRRC37A4P | 0.52 |
| LRRC38 | 0.69 |
| LRRC3B | 0.60 |
| LRRC49 | 0.73 |
| LRRC59 | 0.53 |
| LRRN1 | 0.52 |
| LTB | 0.50 |
| LURAP1L | 0.54 |
| LUZP1 | 0.76 |
| LUZP2 | 0.63 |
| LXN | 0.81 |
| LY6H | 0.52 |
| LYPD1 | 0.55 |
| LYPLA1 | 0.60 |
| LYPLAL1 | 0.52 |
| LYRM9 | 0.67 |
| LYZL4 | 0.51 |
| MACROD2 | 0.58 |
| MADCAM1 | 0.51 |
| MAFG | 0.51 |
| MAGI3 | 0.62 |
| MAMLD1 | 0.50 |
| MAN1A1 | 0.54 |
| MAOA | 0.55 |
| MAOB | 0.74 |
| MAP2K6 | 0.55 |
| MAP3K13 | 0.63 |
| MAP9 | 0.62 |
| MAPK1 | 0.63 |
| MAPK3 | 0.50 |
| MARCKS | 0.62 |
| MARCO | 0.68 |
| MAT2B | 0.54 |
| MBOAT2 | 0.66 |
| MCUB | 0.58 |
| MED11 | 0.52 |
| MEPE | 0.58 |
| MESP1 | 0.54 |
| MET | 0.74 |
| METTL24 | 0.63 |
| METTL7A | 0.52 |
| MGP | 0.50 |
| MGST1 | 0.71 |

| | |
|---|---|
| MGST2 | 0.66 |
| MICA | 0.51 |
| MICU3 | 0.67 |
| MINK1 | 0.52 |
| MIOS | 0.53 |
| MIR29B2CHG | 0.73 |
| MIR31HG | 0.63 |
| MIR600HG | 0.54 |
| MIR99AHG | 0.55 |
| MIRLET7BHG | 0.53 |
| MKNK2 | 0.59 |
| MKX | 0.60 |
| MLC1 | 0.51 |
| MMD | 0.65 |
| MOB1B | 0.56 |
| MORC2-AS1 | 0.51 |
| MOXD1 | 0.67 |
| MPP1 | 0.66 |
| MPP6 | 0.59 |
| MR1 | 0.57 |
| MRAS | 0.67 |
| MREG | 0.57 |
| MRGPRF | 0.61 |
| MROH5 | 0.79 |
| MRPL48 | 0.55 |
| MTA3 | 0.54 |
| MTAP | 0.52 |
| MTCL1 | 0.68 |
| MTFP1 | 0.58 |
| MUCL1 | 0.51 |
| MUM1L1 | 0.55 |
| MYB | 0.55 |
| MYBPC1 | 0.67 |
| MYH7 | 0.63 |
| MYH7B | 0.69 |
| MYL5 | 0.64 |
| MYO15A | 0.57 |
| MYO19 | 0.59 |
| N4BP2 | 0.67 |
| NADK2 | 0.72 |
| NANOS1 | 0.68 |
| NAP1L2 | 0.54 |
| NAPEPLD | 0.56 |
| NAT8L | 0.53 |
| NCAM2 | 0.58 |
| NCAN | 0.61 |
| NCK2 | 0.52 |
| NCOA3 | 0.68 |
| NDRG3 | 0.62 |
| NEB | 0.63 |
| NEBL-AS1 | 0.55 |
| NECAB2 | 0.72 |

| | |
|---|---|
| NEFH | 0.80 |
| NEXN | 0.52 |
| NFIB | 0.57 |
| NFYC | 0.53 |
| NGB | 0.54 |
| NIPAL2 | 0.54 |
| NKAIN2 | 0.51 |
| NKAIN3 | 0.65 |
| NKAIN4 | 0.63 |
| NKX6-3 | 0.53 |
| NOL4 | 0.58 |
| NOV | 0.62 |
| NOXO1 | 0.57 |
| NPPA | 0.50 |
| NPY1R | 0.69 |
| NR1D2 | 0.73 |
| NR2C1 | 0.54 |
| NR2F2 | 0.64 |
| NR3C1 | 0.72 |
| NR3C2 | 0.59 |
| NRAP | 0.51 |
| NRIP2 | 0.62 |
| NSG2 | 0.66 |
| NT5DC3 | 0.53 |
| NT5E | 0.54 |
| NT5M | 0.64 |
| NTSR1 | 0.70 |
| NTSR2 | 0.59 |
| NUDT10 | 0.63 |
| NUDT11 | 0.66 |
| NUDT14 | 0.61 |
| NUPR2 | 0.61 |
| NWD1 | 0.51 |
| OIP5-AS1 | 0.71 |
| ONECUT2 | 0.70 |
| OPHN1 | 0.64 |
| OPRK1 | 0.57 |
| OPRM1 | 0.53 |
| OR2L3 | 0.52 |
| OR2L8 | 0.51 |
| OSBP2 | 0.55 |
| OSBPL6 | 0.61 |
| OVOL2 | 0.57 |
| P2RX6 | 0.59 |
| P2RY1 | 0.51 |
| PACSIN3 | 0.52 |
| PALLD | 0.52 |
| PAMR1 | 0.62 |
| PANX2 | 0.50 |
| PAX8-AS1 | 0.58 |
| PBX4 | 0.60 |
| PBXIP1 | 0.57 |

| | |
|---|---|
| PCBD1 | 0.51 |
| PCDH10 | 0.54 |
| PCDH12 | 0.67 |
| PCDH15 | 0.51 |
| PCDH17 | 0.73 |
| PCDH19 | 0.50 |
| PCDH20 | 0.60 |
| PCED1B | 0.57 |
| PCGF1 | 0.51 |
| PCP4 | 0.71 |
| PCP4L1 | 0.53 |
| PCSK5 | 0.77 |
| PDGFC | 0.57 |
| PDYN | 0.61 |
| PEA15 | 0.75 |
| PELI3 | 0.53 |
| PGAP1 | 0.69 |
| PGM2L1 | 0.54 |
| PGRMC1 | 0.60 |
| PHLDA2 | 0.56 |
| PHYH | 0.63 |
| PI4K2A | 0.51 |
| PID1 | 0.72 |
| PIF1 | 0.55 |
| PKD2L1 | 0.53 |
| PKIA | 0.75 |
| PLCD4 | 0.56 |
| PLCH1 | 0.68 |
| PLEKHH3 | 0.55 |
| PLEKHO2 | 0.52 |
| PLK5 | 0.62 |
| PLPP3 | 0.53 |
| PLPPR3 | 0.59 |
| PLPPR4 | 0.60 |
| PLSCR4 | 0.66 |
| PLTP | 0.57 |
| PLXDC1 | 0.70 |
| PLXNA1 | 0.52 |
| PLXNC1 | 0.66 |
| PNISR | 0.54 |
| PNKD | 0.71 |
| PNLDC1 | 0.55 |
| PNMT | 0.78 |
| PNPLA3 | 0.63 |
| POLE4 | 0.55 |
| POLR2L | 0.52 |
| PON2 | 0.52 |
| PON3 | 0.62 |
| POU6F1 | 0.50 |
| POU6F2 | 0.55 |
| PPARA | 0.54 |
| PPARGC1A | 0.77 |

| | |
|---|---|
| PPEF1 | 0.55 |
| PPIL3 | 0.55 |
| PPM1M | 0.63 |
| PPP2R5C | 0.51 |
| PPP4R4 | 0.72 |
| PPTC7 | 0.69 |
| PRAG1 | 0.50 |
| PRDM16 | 0.53 |
| PRDM2 | 0.54 |
| PRKCD | 0.66 |
| PRKCG | 0.71 |
| PRKG1 | 0.55 |
| PRMT7 | 0.65 |
| PRO1804 | 0.59 |
| PRODH | 0.53 |
| PRRX1 | 0.70 |
| PRSS16 | 0.62 |
| PRSS35 | 0.56 |
| PRUNE2 | 0.52 |
| PSIP1 | 0.59 |
| PSMD12 | 0.64 |
| PSTPIP1 | 0.55 |
| PTCHD1 | 0.54 |
| PTER | 0.54 |
| PTGER3 | 0.76 |
| PTH2R | 0.52 |
| PTPN4 | 0.62 |
| PTPRA | 0.58 |
| PTPRD-AS1 | 0.51 |
| PVALB | 0.80 |
| PXDNL | 0.53 |
| PYDC1 | 0.59 |
| PYGL | 0.59 |
| RAB27B | 0.68 |
| RAB31 | 0.62 |
| RAB37 | 0.69 |
| RAB3C | 0.50 |
| RAD54B | 0.74 |
| RADIL | 0.56 |
| RAMP3 | 0.52 |
| RANBP3L | 0.56 |
| RAP2B | 0.60 |
| RAPGEF4 | 0.63 |
| RARB | 0.62 |
| RASAL1 | 0.57 |
| RASSF4 | 0.57 |
| RAVER2 | 0.74 |
| RBMS1 | 0.58 |
| RBP4 | 0.76 |
| RCAN2 | 0.68 |
| RCN3 | 0.56 |
| RELL2 | 0.68 |

| | |
|---|---|
| RELT | 0.59 |
| RET | 0.61 |
| RFPL2 | 0.57 |
| RFX3 | 0.52 |
| RGS20 | 0.64 |
| RGS22 | 0.52 |
| RHBDL3 | 0.72 |
| RHOBTB2 | 0.58 |
| RHOC | 0.52 |
| RILP | 0.63 |
| RILPL2 | 0.71 |
| RIMKLA | 0.65 |
| RIPOR2 | 0.77 |
| RMDN2 | 0.54 |
| RNF113A | 0.50 |
| ROBO1 | 0.54 |
| RORA | 0.72 |
| RPGR | 0.64 |
| RPP25 | 0.55 |
| RRAGB | 0.51 |
| RRP7A | 0.57 |
| RS1 | 0.62 |
| RSPH9 | 0.77 |
| RSRC1 | 0.54 |
| RTKN2 | 0.62 |
| RTP1 | 0.73 |
| RYR3 | 0.54 |
| S100A13 | 0.56 |
| SAA2 | 0.50 |
| SAP30BP | 0.51 |
| SARAF | 0.51 |
| SAT2 | 0.50 |
| SCAPER | 0.53 |
| SCD5 | 0.52 |
| SCN1A | 0.71 |
| SCN1B | 0.81 |
| SCN3A | 0.64 |
| SCN3B | 0.75 |
| SCN4B | 0.76 |
| SCN9A | 0.76 |
| SCRG1 | 0.58 |
| SCRT1 | 0.69 |
| SDC2 | 0.55 |
| SDSL | 0.68 |
| SECISBP2 | 0.51 |
| SEMA3D | 0.58 |
| SEMA7A | 0.72 |
| SERTAD4 | 0.67 |
| SERTAD4-AS1 | 0.63 |
| SETD7 | 0.60 |
| SEZ6 | 0.51 |
| SH2D1B | 0.51 |

| | |
|---|---|
| SH3BGRL3 | 0.50 |
| SH3RF1 | 0.54 |
| SH3RF2 | 0.52 |
| SHD | 0.78 |
| SHF | 0.55 |
| SHISAL1 | 0.68 |
| SHROOM3 | 0.58 |
| SIRT4 | 0.53 |
| SIX4 | 0.67 |
| SKAP1 | 0.59 |
| SKAP2 | 0.57 |
| SLA | 0.72 |
| SLC15A2 | 0.53 |
| SLC16A2 | 0.63 |
| SLC16A6 | 0.74 |
| SLC16A7 | 0.64 |
| SLC17A6 | 0.74 |
| SLC17A8 | 0.62 |
| SLC19A2 | 0.51 |
| SLC1A3 | 0.55 |
| SLC20A2 | 0.55 |
| SLC22A10 | 0.56 |
| SLC22A18 | 0.51 |
| SLC24A2 | 0.60 |
| SLC25A12 | 0.58 |
| SLC25A18 | 0.56 |
| SLC25A23 | 0.52 |
| SLC25A25 | 0.53 |
| SLC25A37 | 0.58 |
| SLC25A5 | 0.52 |
| SLC30A10 | 0.55 |
| SLC38A1 | 0.60 |
| SLC39A10 | 0.57 |
| SLC39A12 | 0.53 |
| SLC39A13 | 0.58 |
| SLC39A14 | 0.59 |
| SLC47A1 | 0.58 |
| SLC4A4 | 0.58 |
| SLC4A8 | 0.63 |
| SLC7A10 | 0.54 |
| SLC7A11 | 0.57 |
| SLC7A4 | 0.59 |
| SLCO1C1 | 0.50 |
| SLCO4A1 | 0.56 |
| SLF1 | 0.52 |
| SLIT1 | 0.74 |
| SMARCD3 | 0.58 |
| SMIM10L2A | 0.56 |
| SMIM3 | 0.51 |
| SMIM30 | 0.53 |
| SMO | 0.55 |
| SMPX | 0.61 |

| | |
|---|---|
| SNHG1 | 0.51 |
| SNHG14 | 0.56 |
| SNRNP27 | 0.53 |
| SNTA1 | 0.51 |
| SNX24 | 0.51 |
| SNX7 | 0.60 |
| SOCS5 | 0.53 |
| SOHLH1 | 0.60 |
| SORL1 | 0.66 |
| SOWAHA | 0.58 |
| SOX5 | 0.58 |
| SOX9-AS1 | 0.56 |
| SPAG4 | 0.67 |
| SPATA18 | 0.73 |
| SPATA33 | 0.64 |
| SPHK1 | 0.53 |
| SPHKAP | 0.68 |
| SPON2 | 0.64 |
| SPRR2G | 0.51 |
| SPTSSB | 0.70 |
| SRGAP1 | 0.56 |
| SRPK1 | 0.69 |
| SSBP2 | 0.56 |
| SSTR1 | 0.80 |
| ST3GAL6 | 0.69 |
| ST3GAL6-AS1 | 0.66 |
| ST6GALNAC5 | 0.55 |
| ST8SIA1 | 0.50 |
| STAC2 | 0.75 |
| STARD10 | 0.55 |
| STARD5 | 0.63 |
| STK17A | 0.56 |
| STK39 | 0.60 |
| STON2 | 0.57 |
| STRBP | 0.70 |
| STS | 0.56 |
| STUM | 0.72 |
| STX17-AS1 | 0.55 |
| STX19 | 0.54 |
| STX7 | 0.56 |
| STXBP5L | 0.64 |
| SUB1 | 0.55 |
| SULF1 | 0.54 |
| SULF2 | 0.66 |
| SUSD4 | 0.50 |
| SV2C | 0.83 |
| SYCP2 | 0.77 |
| SYN2 | 0.61 |
| SYNE4 | 0.51 |
| SYT10 | 0.53 |
| SYT12 | 0.58 |
| SYT17 | 0.77 |

| | |
|---|---|
| SYT2 | 0.79 |
| SYT6 | 0.51 |
| SYTL2 | 0.50 |
| TAF9B | 0.63 |
| TBC1D24 | 0.51 |
| TDP2 | 0.50 |
| TDRD1 | 0.76 |
| TEAD4 | 0.52 |
| TENM3 | 0.64 |
| TESMIN | 0.70 |
| TEX26 | 0.64 |
| TEX29 | 0.69 |
| TFAM | 0.58 |
| TG | 0.67 |
| TGFBI | 0.64 |
| THAP10 | 0.65 |
| THEMIS | 0.52 |
| THRA | 0.65 |
| TIAM1 | 0.56 |
| TIFA | 0.61 |
| TIMP3 | 0.51 |
| TINCR | 0.63 |
| TLE2 | 0.52 |
| TM2D3 | 0.53 |
| TMEFF2 | 0.67 |
| TMEM132E | 0.57 |
| TMEM150C | 0.53 |
| TMEM158 | 0.53 |
| TMEM159 | 0.58 |
| TMEM161B-AS1 | 0.54 |
| TMEM196 | 0.67 |
| TMEM200A | 0.57 |
| TMEM200B | 0.65 |
| TMEM229B | 0.53 |
| TMEM249 | 0.55 |
| TMEM263 | 0.62 |
| TMEM47 | 0.52 |
| TMEM81 | 0.50 |
| TMOD1 | 0.56 |
| TMPO | 0.54 |
| TMSB10 | 0.53 |
| TMTC1 | 0.61 |
| TMX2 | 0.51 |
| TMX4 | 0.53 |
| TNFRSF11A | 0.62 |
| TNNT1 | 0.62 |
| TNNT2 | 0.52 |
| TNS3 | 0.51 |
| TOM1L1 | 0.55 |
| TP53BP2 | 0.50 |
| TP53I11 | 0.59 |
| TPBG | 0.65 |

| | |
|---|---|
| TPD52 | 0.53 |
| TPTE2P6 | 0.53 |
| TRANK1 | 0.59 |
| TRIL | 0.61 |
| TRIM36 | 0.60 |
| TRIM37 | 0.52 |
| TRIM52 | 0.53 |
| TRIM7 | 0.56 |
| TRMT61B | 0.63 |
| TRMT9B | 0.59 |
| TRPC3 | 0.62 |
| TSHZ1 | 0.65 |
| TSPAN1 | 0.69 |
| TSPAN33 | 0.56 |
| TSPOAP1 | 0.50 |
| TTC21B | 0.55 |
| TTC39A | 0.56 |
| TTC39B | 0.63 |
| TTR | 0.50 |
| TUBB2A | 0.54 |
| TUBE1 | 0.55 |
| TUBGCP5 | 0.51 |
| TUNAR | 0.75 |
| UBE2QL1 | 0.56 |
| UCHL3 | 0.67 |
| UCHL5 | 0.65 |
| UG0898H09 | 0.67 |
| UHMK1 | 0.69 |
| ULK3 | 0.57 |
| UNC5B-AS1 | 0.76 |
| UPP1 | 0.68 |
| UST | 0.50 |
| VAMP1 | 0.77 |
| VAT1L | 0.62 |
| VAV3 | 0.71 |
| VBP1 | 0.52 |
| WDR1 | 0.54 |
| WDR6 | 0.51 |
| WDR66 | 0.68 |
| WDR86 | 0.70 |
| WDR97 | 0.75 |
| WFDC1 | 0.60 |
| WIF1 | 0.63 |
| WNT10B | 0.60 |
| WTIP | 0.57 |
| XKR4 | 0.52 |
| XKR9 | 0.50 |
| XYLT1 | 0.63 |
| YAP1 | 0.59 |
| YPEL1 | 0.71 |
| ZADH2 | 0.74 |
| ZBTB1 | 0.56 |

| | |
|---|---|
| ZBTB8A | 0.59 |
| ZCCHC12 | 0.70 |
| ZCCHC18 | 0.68 |
| ZDHHC2 | 0.53 |
| ZFPM2 | 0.50 |
| ZFYVE9 | 0.52 |
| ZIM2 | 0.56 |
| ZMAT1 | 0.52 |
| ZNF148 | 0.51 |
| ZNF165 | 0.54 |
| ZNF268 | 0.65 |
| ZNF385B | 0.61 |
| ZNF385D | 0.53 |
| ZNF630 | 0.51 |
